# Hedgehog-dependent and Hedgehog-independent roles for Growth Arrest Specific 1 in mammalian kidney morphogenesis

**DOI:** 10.1101/2024.05.02.592197

**Authors:** Nicole E. Franks, Benjamin L. Allen

## Abstract

Growth arrest specific 1 (GAS1) is a key regulator of mammalian embryogenesis, best known for its role in Hedgehog (HH) signaling, but with additional described roles in the FGF, RET, and NOTCH pathways. Previous work indicated a later role for GAS1 in kidney development through FGF pathway modulation. Here, we demonstrate that GAS1 is essential for both mesonephrogenesis and metanephrogenesis– most notably, *Gas1* deletion in mice results in renal agenesis in a genetic background-dependent fashion. Mechanistically, GAS1 promotes mesonephrogenesis in a HH-dependent fashion, performing a unique co-receptor function, while promoting metanephrogenesis in a HH-independent fashion, acting as a putative secreted RET co-receptor. Our data indicate that *Gas1* deletion leads to renal agenesis through a transient reduction in metanephric mesenchyme proliferation– a phenotype that can be rescued by exogenous RET pathway stimulation. Overall, this study indicates that GAS1 contributes to early kidney development through the integration of multiple different signaling pathways.

## Introduction

A long-standing tenet of developmental biology is that proper organogenesis requires cells to integrate a multitude of signals in a spatial- and time-dependent fashion. While previous research efforts have succeeded in identifying individual pathway requirements for organ formation, there remains a general lack of understanding regarding the synthesis of multiple signaling inputs to execute a specific cellular response, and even less understanding of how subsequent communication between cell types drives appropriate tissue development. One emerging theme that addresses the issue of complex organ formation is the utilization of individual proteins that can impact multiple signaling pathways within a cell or across a tissue. Investigating these proteins may help to reveal the mechanisms used to ensure proper organ development.

Growth arrest specific 1 (GAS1) is a multi-functional, cell surface-associated protein that plays essential roles throughout mammalian embryogenesis (Allen et al., 2011; Allen et al., 2007; Biau et al., 2013; Lee et al., 2001; Martinelli and Fan, 2007a, b; Seppala et al., 2007).

*Gas1* was first identified as a gene up-regulated in serum-starved NIH/3T3 fibroblasts (Schneider et al., 1988) and encodes a 45kDa Glycosylphosphatidylinositol (GPI)-anchored protein that shares structural homology with Glial-derived neurotrophic factor (GDNF) receptors (GFRs) (Cabrera et al., 2006). During embryogenesis, GAS1 is broadly expressed, and is best studied as a co-receptor for the Hedgehog (HH) signaling pathway, where it is required for the patterning and growth of multiple tissues (Allen et al., 2011; Allen et al., 2007; Biau et al., 2013; Izzi et al., 2011; Lee et al., 2001; Lee and Fan, 2001; Liu et al., 2001; Martinelli and Fan, 2007a).

Initial studies described an antagonistic role for GAS1 in HH signaling in early tooth (Lee et al., 2001) and somite development (Cobourne et al., 2004; Ohazama et al., 2009).

However, more recent work in mouse embryos demonstrated a role for *Gas1* in promoting HH pathway activity, where *Gas1* deletion results in HH-dependent defects in craniofacial development, digit specification, and ventral neural tube patterning (Allen et al., 2011; Allen et al., 2007; Martinelli and Fan, 2007a). Notably, GAS1 acts in coordination with two additional cell surface proteins, CAM-related/downregulated by oncogenes (CDON), and brother of CDON (BOC), which together are required for HH signal transduction (Allen et al., 2011; Allen et al., 2007; Echevarria-Andino and Allen, 2020; Echevarria-Andino et al., 2022; Izzi et al., 2011; Martinelli and Fan, 2007a).

In addition to its role in HH signaling, GAS1 has ascribed functions in the regulation of other signaling pathways. In particular, the structural similarity of GAS1 to GFRs led to work demonstrating physical interactions between GAS1 and the receptor tyrosine kinase RET (Cabrera et al., 2006). Further, GAS1 has been described as a negative regulator of RET signaling in cultured cells (Biau et al., 2013; Cabrera et al., 2006; Li et al., 2019; Lopez-Ramirez et al., 2008). More recent work has also identified in vivo roles for GAS1 in modulating RET pathway activation during gastrointestinal development (Biau et al., 2013) and in skeletal muscle stem cell self-renewal (Li et al., 2019). Moreover, GAS1 has described functions as a regulator of NOTCH-dependent SHH signaling in forebrain morphogenesis (Marczenke et al., 2021).

In the developing kidney, it has been proposed that GAS1 functions independently of its role in HH-signaling. Specifically, *Gas1* deletion in mice on a mixed genetic background results in renal hypoplasia at later developmental stages (i.e., after embryonic day E15.5) (Kann et al., 2015). It was suggested that *Gas1* modulation of FGF signaling during kidney morphogenesis is responsible for the renal hypoplasia phenotypes observed in *Gas1* mutants (Kann et al., 2015). However, RET signaling has essential and well-studied roles in kidney morphogenesis, while HH signaling also has demonstrated contributions to multiple aspects of both mesonephric and metanephric development (Bohnenpoll et al., 2017; Cain et al., 2009; Chung et al., 2024; Murashima et al., 2014; Rowan et al., 2018; Yu et al., 2002). To date, potential roles for GAS1 in the regulation of these signaling pathways during kidney development remain unexplored.

Here we investigated the contributions of GAS1 to kidney morphogenesis in mouse embryonic development. Surprisingly, we found that *Gas1* deletion on a congenic C57BL6/J background results in renal agenesis, a phenotype previously undescribed in *Gas1* mutant animals. Further, we show that these phenotypes are genetic background-dependent, as *Gas1* mutant animals on a congenic 129S4/SvJaeJ background do not display renal agenesis but instead display renal hypoplasia as early as E12.5; again, a previously unreported phenotype. Further, we found that *Gas1* mutants across multiple genetic backgrounds display ectopic caudal mesonephric tubules and decreased HH target gene expression, a phenotype consistent with reduced HH pathway activity. Surprisingly, this phenotype was specific to *Gas1*, as individual or combined deletion of *Cdon* or *Boc* did not result in any overt kidney phenotypes.

In *Gas1* mutant metanephroi, we observed defects in mesenchyme proliferation and survival in early kidney morphogenesis. Further, *Gas1* deletion contributes to decreased expression of downstream RET transcriptional targets in the metanephros. Mechanistically, our data suggest that GAS1 is secreted from the kidney mesenchyme and can physically interact with RET in the adjacent renal epithelium. Ex vivo kidney explant cultures demonstrate that the branching morphogenesis defects observed in *Gas1* mutant kidneys can be rescued by GDNF-mediated RET pathway activation. Together, these data indicate that GAS1 is a multi-functional regulator of kidney development, employing both HH-dependent and HH-independent mechanisms to drive proper mesonephros formation and kidney branching morphogenesis.

## Results

### *Gas1* is an essential regulator of early kidney development

While previous work identified a role for *Gas1* in later stages of kidney morphogenesis (i.e., after E15.5 in mouse; (Kann et al., 2015) our investigations of *Gas1* function in embryogenesis led to a fundamentally different observation. Specifically, analysis of wildtype (Fig. 1A), *Gas1^+/-^*(Fig. 1B), and *Gas1^-/-^* (Fig. 1C-E) embryos at E15.5 revealed a range of developmental defects in kidney morphogenesis in *Gas1* mutants on a C57BL/6J background, from renal hypoplasia (Fig. 1C) to unilateral renal agenesis (Fig. 1D) to bilateral renal agenesis (Fig. 1E). Notably, the bilateral renal agenesis observed in a subset of *Gas1* mutants phenocopies that observed in *Ret* mutant embryos (Schuchardt et al., 1994); Fig. 1F). Quantitation of these phenotypes indicated normal kidney development in 100% of wildtype and *Gas1^+/-^* embryos, while *Gas1^-/-^* embryos most frequently display unilateral renal agenesis (64%); renal hypoplasia (14%) and bilateral renal agenesis (22%) occur less frequently (Fig. 1G).

**Figure 1.**
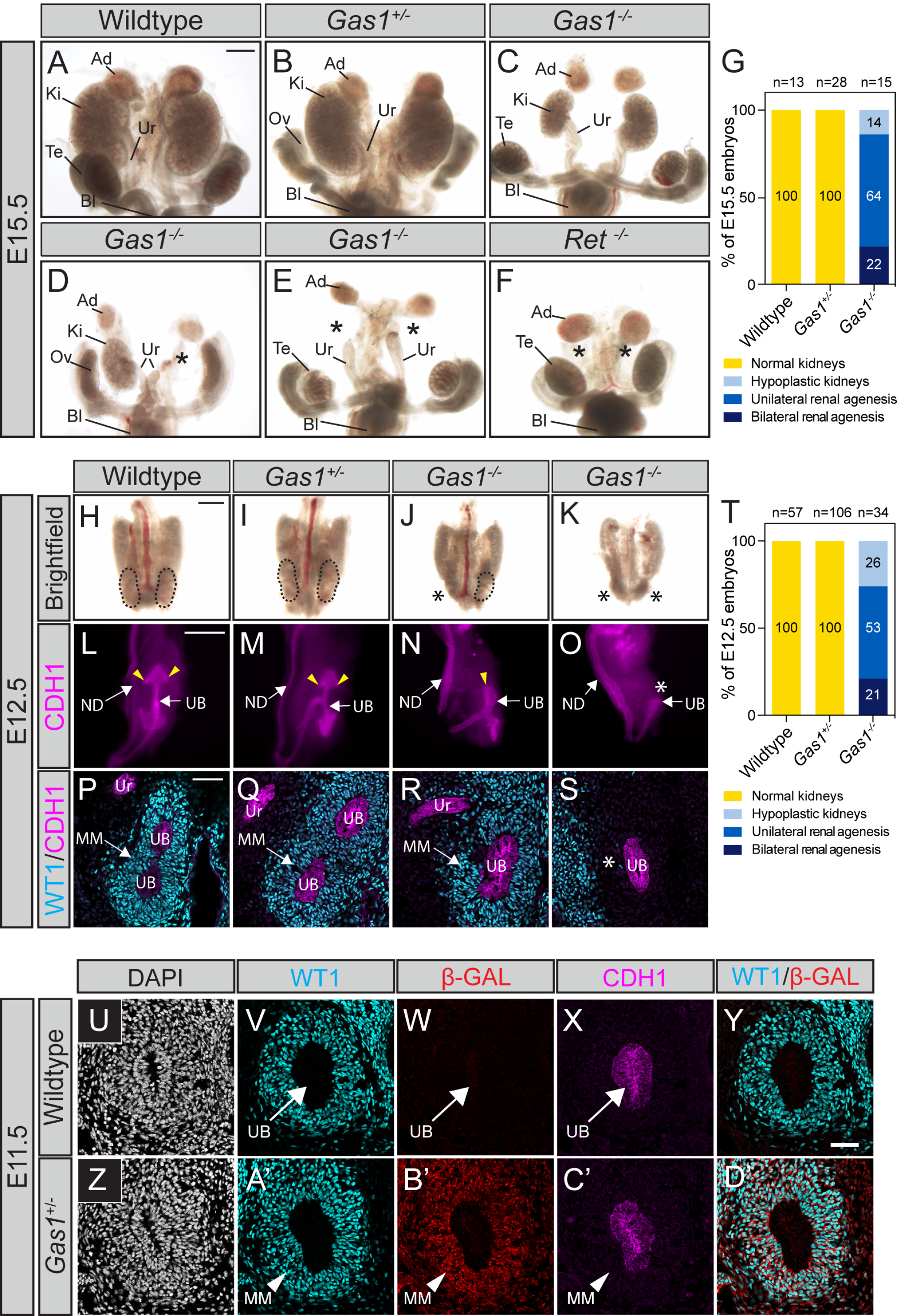
*Gas1^-/-^* embryos on a C57BL/6J background display renal hypoplasia and agenesis. (A-F) Whole-mount brightfield images of E15.5 wildtype (A), *Gas1^+/-^* (B), *Gas1^-/-^* (C-E) and *Ret^-/-^* (F) mouse embryonic kidneys. (G) Quantitation of the frequency of renal defects in wildtype (n=13), *Gas1^+/-^* (n=28), and *Gas1^-/-^* (n=15) E15.5 embryos. Embryos were classified into the following categories: normal kidneys, hypoplastic kidneys, unilateral renal agenesis, or bilateral renal agenesis. (H-K) Whole-mount brightfield images of E12.5 (52-56 somites) wildtype (H), *Gas1^+/-^* (I), and *Gas1^-/-^* (J,K) mouse kidneys (marked by dotted lines). (L-O) Whole-mount immunofluorescent detection of E-Cadherin (CDH1) in E12.5 wildtype (L), *Gas1^+/-^* (M) and *Gas1^-/-^* (N,O) kidneys. Yellow arrowheads identify ureteric bud epithelial branching events (L,M,N). (P-S) Antibody detection of CDH1 and WT1 in sections from E12.5 wildtype (P), *Gas1^+/-^* (Q), and *Gas1^-/-^* (R,S) kidneys. (T) Quantitation of the frequency of renal defects in E12.5 wildtype (n=57), *Gas1^+/-^* (n=106), and *Gas1^-/-^* (n=34) embryos. (U-D’) Immunofluorescent confocal microscope images of coronal sections of metanephros in E11.5 wildtype and *Gas1^lacZ/+^* mouse embryos. Nuclei are labeled with DAPI (U,Z). Antibody detection of WT1 (cyan; V,A’), β-galactosidase (β-GAL, red; W,B’), CDH1 (magenta; X,C’). Merged WT1/β -GAL images are shown (Y,D’). Asterisks denote renal agenesis (D-F,J,K,O,S). Scale bars (A, H) 500μm, (L) 200μm, (P) 50μm. Abbreviations: ureter (Ur), adrenal gland (Ad), kidney (Ki), testis (Te), ovary (Ov), ureteric bud (UB), nephric duct (ND), bladder (Bl), metanephric mesenchyme (MM).

To investigate the onset of these phenotypes, we collected *Gas1* mutant embryos at E12.5, a developmental stage that reflects early branching events of the ureteric bud (UB) epithelium, a structure that gives rise to the renal collecting duct system (Fig. 1H-T). Again, wildtype (Fig. 1H) and *Gas1^+/-^* (Fig. 1I) embryos display normal metanephros development; however, *Gas1^-/-^* embryos are perturbed even at this early stage of renal development (Fig. 1J, K), with hypoplasia (26%), unilateral renal agenesis (53%) and bilateral renal agenesis (21%) occurring at similar frequencies to those observed at E15.5 (cf. Fig. 1G and 1T).

Immunofluorescent antibody detection of E-Cadherin (CDH1) confirmed initial UB branching events in wildtype (Fig. 1L) and *Gas1^+/-^* (Fig. 1M) embryos, while branching of the ureteric bud was observed in hypoplastic *Gas1^-/-^* kidneys (Fig. 1N), but not in *Gas1* mutants lacking kidneys (Fig. 1O). Notably, ureteric bud branching is delayed in hypoplastic *Gas1^-/-^* kidneys compared with wildtype or *Gas1^+/-^* kidneys (cf. Fig. 1L-N). Given the importance of epithelial-mesenchymal signaling in metanephros branching morphogenesis (Combes et al., 2015; Costantini and Kopan, 2010; Little and McMahon, 2012), we used antibody detection of Wilms Tumor 1 (WT1) to assess the metanephric mesenchyme (MM) in wildtype and *Gas1* mutant embryos at E12.5 (Fig. 1P-S). Strikingly, WT1 staining indicates a qualitative difference between hypoplastic *Gas1* mutant kidneys, where WT1+ cells are observed (Fig. 1R), and *Gas1^-/-^* embryos displaying renal agenesis, which lack WT1+ cells in the immediate vicinity of the ureteric bud (Fig. 1S). To assess whether *Gas1* is expressed during these initial branching events of the ureteric bud, we utilized a *Gas1^lacZ^* reporter allele (Martinelli and Fan, 2007a) to read out *Gas1* expression at the onset of kidney branching morphogenesis. At the first ureteric bud branching event (E11.5), we observed *Gas1* expression in WT1+ MM, whereas *Gas1* was not detected in the adjacent CDH1+ UB epithelium (Fig. 1U-D’). We further confirmed this MM-restricted *Gas1* expression by co-staining with SIX2, a marker specific to nephron progenitor cells (NPCs; Fig. S1A-J). Notably, we observed co-expression of WT1 and *Gas1* in the MM as early as E10.5, a stage that reflects initial specification of the metanephric mesenchyme (Fig. S1K-T).

*Gas1* mutant embryos display genetic background-dependent phenotypes in other tissues during development (Allen et al., 2007; Echevarria-Andino and Allen, 2020; Echevarria-Andino et al., 2022; Martinelli and Fan, 2007a; Seppala et al., 2007; Seppala et al., 2014). Thus, we collected kidneys from wildtype (Fig. S2A,D*) Gas1^+/-^* (Fig. S2B,E) and *Gas1^-/-^* (Fig. S2C,F) mouse embryos maintained on a congenic 129S4/SvJaeJ background to investigate kidney morphogenesis in these animals. At E15.5, we observed normal kidney development in wildtype (Fig. S2A) and *Gas1^+/-^* (Fig. S2B) embryos. In contrast to the variable renal defects observed in *Gas1* mutants on a C57BL6/J genetic background, all *Gas1^-/-^*embryos maintained on a congenic 129S4/SvJaeJ background exhibit bilateral renal hypoplasia (Fig. S2C) which was evident as early as E12.5 (Fig. S2F,G). To confirm that renal hypoplasia is not a result of decreased embryo size in *Gas1* mutants, we measured kidney area and normalized to the crown-rump length in wildtype, *Gas1^+/-^*and *Gas1^-/-^* embryos maintained on congenic 129S4/SvJaeJ (Fig. S2H) and C57BL6/J (Fig. S2I) backgrounds. Notably, this quantitation revealed similar reductions in kidney size in *Gas1* mutants when compared with wildtype littermates (39.5% decrease, 129S4/SvJaeJ, 32.7% decrease, C57BL6/J). These data demonstrate that *Gas1* is required for proper development of the kidney, and that loss of *Gas1* contributes to variable renal defects in a genetic background-dependent fashion.

### Specific HH-dependent role for *Gas1* in mesonephrogenesis

Previous work identified a role for HH signaling in the development of the mesonephros (Murashima et al., 2014), a structure that precedes metanephros development (Fig. 2A). Given the known role of *Gas1* in HH signaling (Allen et al., 2011; Allen et al., 2007; Cobourne et al., 2004; Izzi et al., 2011; Lee et al., 2001; Ohazama et al., 2009; Seppala et al., 2007; Seppala et al., 2022) and its expression at the onset of metanephros morphogenesis, we investigated a potential earlier contribution of GAS1 to mesonephros development, along with the HH co-receptors CDON and BOC. We utilized *Gas1^lacZ^, Cdon^lacZ^* and *Boc^hPLAP^* reporter alleles (Cole and Krauss, 2003; Martinelli and Fan, 2007a; Zhang et al., 2011) to define HH co-receptor expression in the developing mouse mesonephros (Fig. 2B-E, Fig. S3A-L). At E12.5, all three co-receptors are expressed in the mesonephros (Fig. 2C-E; Fig S3D,F,J,L); *Gas1* (Fig. 2C) and *Boc* (Fig. 2D; Fig. S3D,F) are broadly expressed in the mesenchyme, whereas *Cdon* expression is more restricted in this region (Fig. 2E; Fig. S3J,L). *Boc* and *Cdon* expression, but not *Gas1,* is present in the mesonephric tubules and nephric duct epithelium (Fig. S3D,F,J,L). We also examined HH co-receptor expression in the metanephros in E12.5 embryos (Fig. 2B-E; Fig. S3B,H), which shows MM-restricted *Gas1* expression, whereas *Boc* is broadly expressed in the UB and MM, and lower levels of *Cdon* are observed in these areas (Fig. 2C,D,E; Fig. S3B,H). Overall, these data demonstrate partially overlapping, but not identical expression patterns for the HH co-receptors in early kidney development.

**Figure 2.**
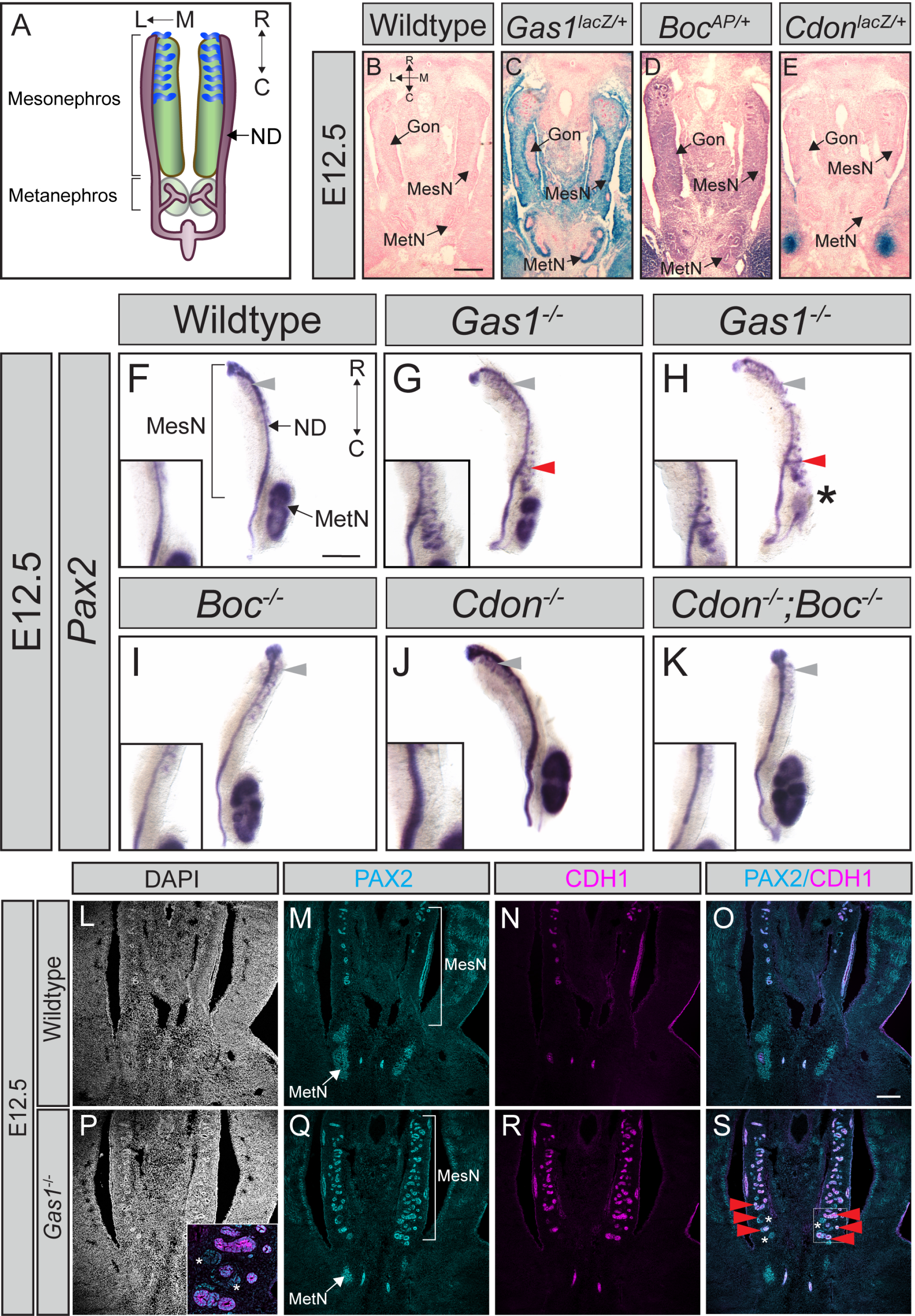
*Gas1* mutants on a congenic C57BL6/J background display ectopic caudal mesonephric tubules. (A) Schematic of wildtype E12.5 mouse mesonephros and metanephros (green, mesenchyme; purple, epithelium; blue, mesonephric tubules). L, lateral; M, medial; R, rostral; C, caudal; ND, nephric duct. (B-E) X-gal (*Gas1, Cdon)* or alkaline phosphatase (AP; *Boc)* staining in mesonephros coronal sections in wildtype (B), *Gas1^lacZ/+^*(C), *Boc^AP/+^* (D) and *Cdon^lacZ/+^*(E) E12.5 embryos. (F-K) Whole-mount *in situ* hybridization for *Pax2* in wildtype (F), *Gas1^-/-^* (G,H), *Boc^-/-^* (I), *Cdon^-/-^* (J) and *Cdon^-/-^;Boc^-/-^*(K) E12.5 mesonephros and metanephros. Insets show a magnified image of the caudal mesonephros. Gray arrowheads indicate rostral mesonephric tubules. Red arrowheads identify ectopic caudal mesonephric tubules (G,H). Asterisks denote renal agenesis (H). (L-S) Immunofluorescent detection of DAPI (gray; L,P), PAX2 (cyan; M,Q) and CDH1 (N,R) in coronal sections of E12.5 wildtype (L-O) and *Gas1^-/-^* (P-S) metanephros and mesonephros. Red arrowheads (S) denote ectopic mesonephric tubules. White asterisks (inset in P; S) mark PAX2+CDH1- ectopic structures in the caudal mesonephros. Scale bars (B,F) 250μm, (O) 100μm. Abbreviations: mesonephros (MesN), metanephros (MetN), gonad (Gon), nephric duct (ND).

To investigate potential contributions of *Gas1*, *Cdon*, and *Boc* to mesonephros development, we collected E12.5 wildtype, *Gas1^-/-^, Boc^-/-^,* and *Cdon^-/-^* mutant embryos and performed whole-mount *in situ* hybridization for *Pax2,* a marker expressed in the nephric duct, ureteric epithelium and metanephric mesenchyme (Dressler et al., 1990)(Fig. 2F-J). Given functional redundancy between CDON and BOC in HH signal transduction, we also generated *Cdon;Boc* double mutant embryos (Fig 2K). In all embryos, *Pax2* expression is detected in the nephric duct and the metanephros. Notably, in a subset of *Gas1* mutant embryos, *Pax2* expression in the MM is reduced, consistent with the variable renal defects observed in these animals (Fig. 2H; cf. Fig. 1J,K). *Pax2* expression is also present in the rostral mesonephric tubules in all embryos (Fig. 2F-K; gray arrowheads). Strikingly, in *Gas1* mutants we observed ectopic, *Pax2*-expressing mesonephric tubules in the caudal mesonephros (Fig. 2G,H; red arrowheads), phenocopying *Shh* conditional knockout mouse embryos (Murashima et al., 2014). In addition, these ectopic *Pax2*+ mesonephric tubules were present in all *Gas1^-/-^* embryos regardless of the degree of agenesis in the metanephros (Fig. 2G,H), a phenotype confirmed by section immunofluorescence for CDH1 and PAX2 in E12.5 wildtype and *Gas1^-/-^*mesonephros sections (Fig. 2L-S). Analysis of *Gas1*^-/-^embryos maintained on a 129S4/SvJaeJ congenic background (Fig. S4A-E) showed similar, but not identical phenotypes to *Gas1* mutants on a C57BL6/J congenic background (Fig. S4G,H). Specifically, *Gas1* mutants on a 129S4/SvJaeJ background display ectopic mesonephric tubules that are not fused to the nephric duct (ND; Fig S4D; red arrowheads), while *Gas1* mutants on a C57BL6/J background display ectopic mesonephric tubules that are attached to the ND (Fig S4G; red arrowheads). In contrast to *Gas1* deletion, individual or combined loss of *Cdon* or *Boc* results in normal mesonephrogenesis (Fig. 2I-K). Notably, deletion of these co-receptors also results in normal metanephros development in *Boc^-/-^* (Fig. S3N), *Cdon^-/-^* (Fig. S3O) and *Cdon^-/-^, Boc^-/-^* (Fig. S3P) embryos maintained on a C57BL6/J congenic background compared with wildtype animals (Fig. S3M,Q). These data suggest that, in contrast to *Gas1*, *Cdon* and *Boc* do not contribute to kidney development.

Given the similar mesonephric abnormalities observed in our *Gas1* mutants and the previously reported *Shh* conditional mutant embryos (Murashima et al., 2014), we examined whether *Gas1* deletion correlates with decreased HH pathway activation in the mesonephros. We collected E12.5 wildtype and *Gas1^-/-^,* along with *Boc^-/-^* and *Cdon^-/-^;Boc*^-/-^ mesonephroi, and performed whole-mount and section *in situ* hybridization (WISH/SISH) for *Gli1,* a direct transcriptional target of the HH pathway (Dai et al., 1999) (Fig 3A-D,H). In all embryos, *Gli1* is detected in the gonads and rostral mesonephros (Fig. 3A-D). Further, we identified robust *Gli1* expression in the mesonephric mesenchyme adjacent to the nephric duct epithelium in wildtype, *Boc^-/-^* and *Cdon^-/-^;Boc^-/-^* embryos (Fig. 3A,C,D,E) . In contrast, *Gas1* mutants revealed a qualitative reduction in *Gli1* expression in the caudal mesonephros, consistent with ectopic caudal mesonephric tubules observed in these embryos (Fig. 3B,H). To quantify changes in HH signaling, we performed section fluorescent *in situ* hybridization (FISH) for *Gli1* expression. Importantly, *Gli1* expression is not detected in E12.5 *Gli1^-/-^* mesonephroi sections compared to wildtype, confirming probe specificity (Fig S5A-H). FISH for *Gli1* in E12.5 wildtype (Fig. 3F,G) and *Gas1^-/-^* (Fig. 3I,J) caudal mesonephros sections revealed a significant reduction in *Gli1* expression in the mesenchyme surrounding the nephric duct in *Gas1* mutant embryos (Fig. 3K). Nephric duct epithelial expression of *Shh* is not significantly altered compared with wildtype embryos (Fig. 5SI-Q), suggesting that reduced mesenchymal *Gli1* expression near the ND in *Gas1* mutants is not due to decreased levels of SHH ligand. Notably, abnormal *Shh* expression is detected in a subset of ectopic mesonephric structures in *Gas1^-/-^* embryos (indicated by red arrowheads; Fig. S5N,P).

**Figure 3.**
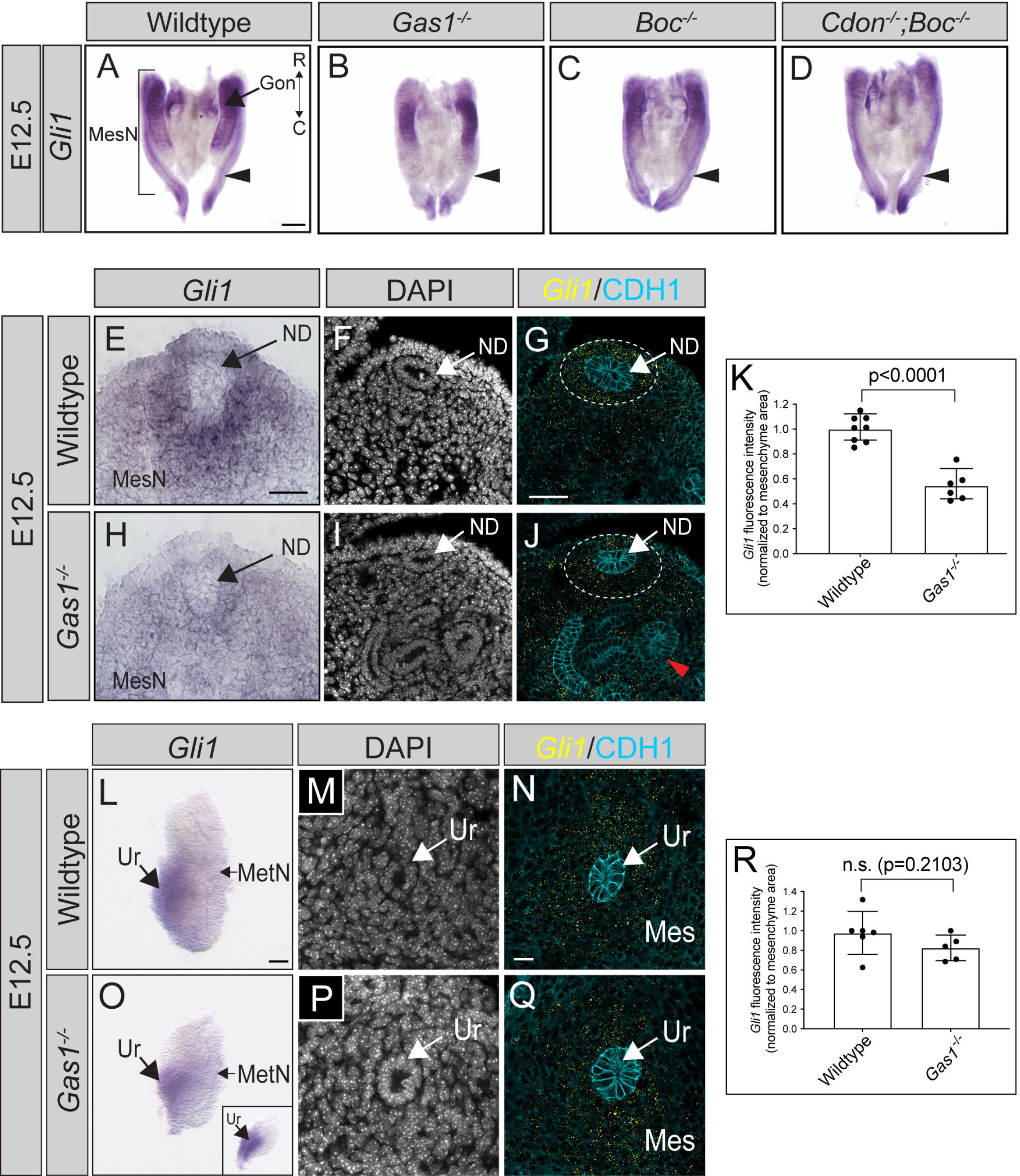
Mesonephric-specific reduction in *Gli1* expression in *Gas1^-/-^* kidneys. (A-D) Whole-mount *in situ* hybridization for *Gli1* in E12.5 wildtype (A), *Gas1^-/-^* (B), *Boc^-/-^* (C) and *Cdon^-/-^* ;*Boc^-/-^* (D) mesonephros. R, rostral; C, caudal; Gon, gonad; MesN, mesonephros. (E,H) Section *in situ* hybridization for *Gli1* in the caudal region of E12.5 wildtype (E) and *Gas1* mutant (H) mesonephros. (F,G,I,J) Representative FISH images for *Gli1* (yellow) with CDH1 (cyan) antibody detection in wildtype (G) and *Gas1^-/-^* (J) transverse caudal mesonephros sections. Nuclei are labeled with DAPI (F,I). ND, nephric duct. (K) Quantitation of *Gli1* fluorescence signal intensity normalized to mesenchyme area in wildtype (n= 8) and *Gas1* mutant (n=6) transverse caudal mesonephros sections. Dotted lines (G,J) represent the mesenchymal area used for normalization of *Gli1* quantitation in K. Red arrowhead denotes ectopic mesonephric tubules (J). (L,O) Whole-mount *in situ* hybridization for *Gli1* in E12.5 wildtype (L), *Gas1^-/-^* hypoplastic kidneys (O) and *Gas1^-/-^* kidneys that have not undergone branching morphogenesis (O, inset). (M,N,P,Q) *Gli1* (yellow) FISH and CDH1 (cyan) protein staining in wildtype (N) and *Gas1* mutant (Q) transverse ureteric mesenchyme sections with DAPI labeling nuclei (M,P). (R) Quantitation of *Gli1* fluorescence signal intensity (normalized to mesenchyme area) in E12.5 wildtype (n=6) and *Gas1^-/-^* (n=5) ureteric mesenchyme. For quantitation (K,R), each data point (n) represents an average of at least 3 sections/kidney. Scale bars (A) 250μm; (E,G,N) 50μm; (L) 100μm. P-values were determined by a two-tailed Student’s t-test: non-significant: p>0.05 and significant: p≤0.05.

HH signaling is required for proper growth and differentiation of the ureteric mesenchyme during metanephros development (Bohnenpoll et al., 2017; Yu et al., 2002). Thus, we analyzed *Gli1* expression in the metanephros in E12.5 wildtype (Fig. 3L-N,R) and *Gas1^-/-^* embryos (Fig. 3O-Q,R) to assess whether *Gas1* contributes to HH signaling in the metanephros. By WISH and FISH analysis, we found that *Gli1* expression is enriched in the ureteral regions in E12.5 wildtype kidneys (Fig. 3L,M,N). In *Gas1^-/-^* mutants, *Gli1* expression is maintained in *Gas1^-/-^* kidneys that have undergone early UB branching morphogenesis or have failed to branch (Fig. 3O, hypoplastic kidney; inset, agenic kidney, Fig. 3P,Q). Quantitation of *Gli1* FISH in E12.5 ureteric mesenchyme sections confirmed no significant changes in *Gli1* expression in *Gas1^-/-^* embryos compared with wildtype animals (Fig. 3R). These data suggest that GAS1 plays a specific, HH-dependent role in mesonephros morphogenesis, while functioning independent of the HH pathway to drive metanephros development.

### *Gas1* promotes proliferation and RET signaling in metanephrogenesis

To investigate potential mechanisms contributing to renal hypoplasia and agenesis in *Gas1* mutant mice, we collected wildtype and *Gas1^-/-^*metanephroi and performed section immunofluorescence to analyze MM (WT1+ or SIX2+) and UB (CDH1+) proliferation or apoptosis by antibody detection of phospho-histone H3 (PHH3) and cleaved caspase-3 (CC3), respectively (Fig 4A-W). At the onset of kidney development (E10.5), the UB and MM is present at the hindlimb level in both wildtype and *Gas1^-/-^*metanephroi embryos, and quantitation of the total number of MM (WT1+CDH1-) cells revealed no significant differences upon *Gas1* loss at this stage (Fig. 4G). However, the number of MM cells undergoing mitosis (PHH3+WT1+CDH1-) is significantly reduced in *Gas1* mutants compared to wildtype animals (Fig. 4H), whereas the percentage of proliferating UB cells (PHH3+CDH1+WT1) remains unchanged (Fig. 4I). In contrast, we did not detect significant changes in MM or UB proliferation at the first UB branching event (E11.5, Fig. 4Q,R); however, the metanephros area is reduced in *Gas1* mutants compared with wildtype animals (Fig. 4J,M), consistent with a transient decrease in MM cell proliferation at an earlier developmental stage (i.e., E10.5). This observation was confirmed by quantitation of total WT1+ MM cells, where we detected approximately 50% fewer MM cells in *Gas1^-/-^* metanephroi at E11.5 compared with wildtype animals (Fig. 4P). At these earliest stages of metanephros development, few CC3+ cells were observed in wildtype (Fig. 4C,L) and *Gas1^-/-^* (Fig. 4F,O) metanephroi. In contrast, we detected a large number of apoptotic MM cells (CC3+SIX2+) that fail to surround the UB in a subset of E12.5 *Gas1* mutant metanephros, indicative of variable renal agenesis in these animals (Fig. 4S-W). Quantitation of CC3+ puncta confirmed no significant changes in the number of apoptotic cells in hypoplastic *Gas1^-/-^* kidneys compared to wildtype embryos, whereas a subset *Gas1* mutant kidneys show increased cell death (Fig. 4W). These results suggest that *Gas1* transiently promotes metanephric mesenchyme proliferation in early kidney development, and that the loss of *Gas1* contributes to MM death in a subset of *Gas1^-/-^* kidneys, which subsequently results in renal agenesis.

**Figure 4.**
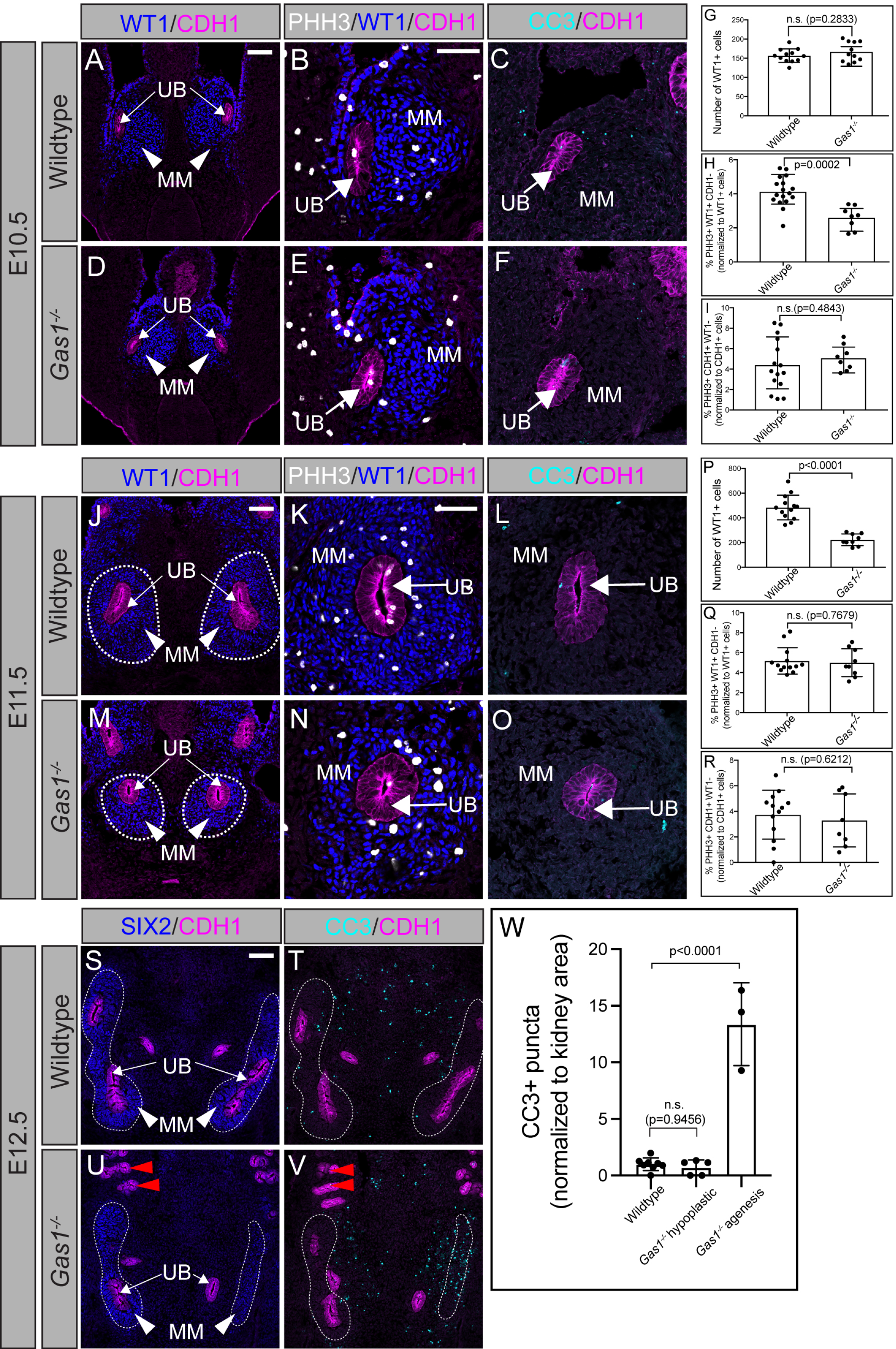
Transient reduction in metanephric mesenchyme proliferation in *Gas1* mutants. (A-F) Coronal section immunofluorescent images of E10.5 wildtype and *Gas1^-/-^*metanephros. Antibody detection of WT1 (blue; A,B,D,E), CDH1 (magenta; A-F), phospho-histone H3 (PHH3, gray; B,E) and cleaved caspase-3 (CC3, cyan; C,F). (G) Quantitation of the total number of WT1+ cells in E10.5 wildtype (n=12) and *Gas1^-/-^* (n=11) metanephros. (H) Quantitation of PHH3+WT1+CDH1- metanephric mesenchyme (MM) cells normalized to the total number of WT1+ cells in coronal sections from E10.5 wildtype (n=17) and *Gas1^-/-^* (n=8) metanephros. (I) Quantitation of PHH3+CDH1+WT1- ureteric bud (UB) cells normalized to the total number of CDH1+ cells in coronal sections from E10.5 wildtype (n=15) and *Gas1^-/-^* (n=8) metanephros. (J-O). Coronal section immunofluorescent images of E11.5 wildtype (J-L) and *Gas1^-/-^* (M-O) metanephros. Antibody detection of WT1 (blue; J,K,M,N), CDH1 (magenta J-O), phospho-histone H3 (PHH3, gray; K,N) and cleaved caspase-3 (CC3, cyan; L,O). (P) Quantitation of the total number of WT1+ cells from E11.5 wildtype (n=13) and *Gas1^-/-^* (n=9) metanephros. (Q) Quantitation of PHH3+WT1+CDH1- metanephric mesenchyme (MM) cells normalized to the total number of WT1+ cells in coronal sections from E11.5 wildtype. (n=13) and *Gas1^-/-^* (n=9) metanephros. (R) Quantitation of PHH3+CDH1+WT1- ureteric bud (UB) cells normalized to the total number of CDH1+ cells in coronal sections from E11.5 wildtype (n=13) and *Gas1^-/-^* (n=8) metanephros. (S-V) Immunofluorescence detection of SIX2 (blue; S,U) and CC3 (T,V) co-stained with CDH1 (magenta; S-V) in serial coronal metanephros sections from E12.5 wildtype and *Gas1*^-/-^ embryos. Red arrowheads mark ectopic caudal mesonephric tubules (U,V). (W) Quantitation of the total number of CC3+ puncta normalized to kidney area in E12.5 wildtype (n=8) and *Gas1^-/-^* hypoplastic (n=5) and agenic (n=3) metanephros. Dotted lines in (S-V) represent the metanephros area used for quantitation. For cell quantitation (G-I,P-R,W), each data point (n) represents the average of at least 2 sections/kidney. Scale bars (A,B,J,K,S), 50μm. P-values were determined by a two-tailed Student’s t-test (G-I, P-R) or an ordinary one-way ANOVA with Tukey’s multiple comparison test (W): non-significant: p>0.05 and significant: p≤0.05.

As shown previously (Fig. 1F), *Gas1*^-/-^ embryos display kidney defects resembling those observed in embryos lacking GDNF/RET signaling (Schuchardt et al., 1994). Therefore, we collected wildtype and *Gas1^-/-^* embryos at the onset of kidney morphogenesis to analyze the expression of several key RET pathway components (Fig. 5A-U;Fig. S6A-Q). Specifically, we collected E10.5 wildtype and *Gas1* mutant metanephroi sections and performed RET antibody detection coupled with FISH for *Gdnf*, which encodes a secreted ligand for RET and the GDNF receptors α (GFRα1-4) (Durbec et al., 1996; Jing et al., 1996; Treanor et al., 1996; Trupp et al., 1996), all components necessary for kidney formation (Cacalano et al., 1998; Moore et al., 1996; Pichel et al., 1996a, b; Sanchez et al., 1996; Schuchardt et al., 1994; Schuchardt et al., 1996) (Fig. 5A-F). Consistent with previous reports (Kume et al., 2000), we observed *Gdnf* expression in wildtype MM at the onset of metanephros development (Fig. 5A-C). Compared with wildtype embryos, *Gdnf* expression is significantly reduced upon loss of *Gas1*, and notably, to a similar degree in all *Gas1^-/-^* metanephroi analyzed (Fig. 5A-G). Further, we analyzed the expression of two cell-surface receptors of the RET pathway, *Ret* and *Gfra1,* in wildtype and *Gas1^-/-^* metanephroi (Fig. S6A-P). In wildtype embryos, we detected *Ret* expression in the nephric duct epithelium prior to UB outgrowth (Fig. S6A) and in the UB at the first branching event (Fig. S6C). Further, *Gfra1* is highly expressed in the UB at E10.5 and E11.0, whereas weak expression is observed in the adjacent MM at E11.0 in wildtype embryos (Fig. S6E,G). In *Gas1* mutants, the expression domain of *Ret* and *Gfra1* confirm delayed UB branching morphogenesis compared to wildtype animals (Fig. S6D,H). However, we did not detect any overt changes in the levels of *Ret* or *Gfra1* expression in these embryos (Fig. S6B,D,F,H), which was confirmed by section antibody detection of RET and GFRA1 in the UB of wildtype, and *Gas1* mutants with renal hypoplasia or agenesis (Fig. S6I-P; agenic, inset). Given the maintained expression of *Ret* and *Gfra1*, and similar reductions in *Gdnf*, these data suggest that additional factors contribute to the variable renal agenesis phenotypes observed in *Gas1* mutant embryos.

**Figure 5.**
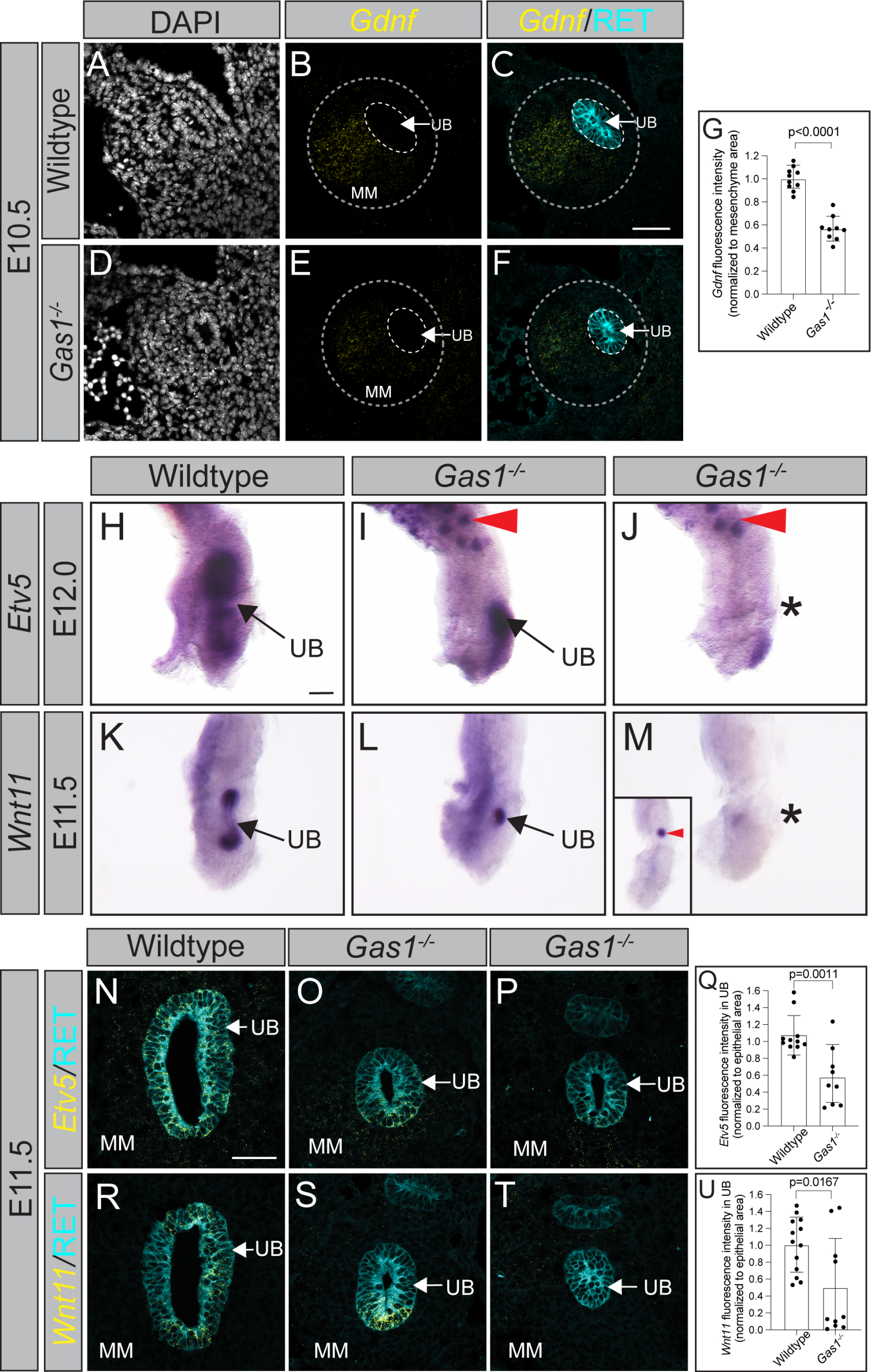
RET signaling is decreased in *Gas1* mutant metanephros. (A-F) FISH for *Gdnf* expression (yellow, B,C,E,F), co-stained with RET (cyan; C,F) antibody detection in E10.5 metanephros coronal sections from wildtype (A-C) and *Gas1^-/-^* (D-F) embryos. DAPI labels nuclei (A,D). (G) Quantitation of *Gdnf* fluorescence signal intensity in E10.5 wildtype (n=10) and *Gas1^-/-^* (n=9) metanephros sections (3 sections/kidney) normalized to metanephric mesenchyme (MM) area. (H-J) Whole-mount *in situ* hybridization of *Etv5* in E12.0 wildtype (H) and *Gas1^-/-^*(I,J) kidneys. Red arrowheads (I,J) indicate ectopic *Etv5-*expressing mesonephric tubules. (K-M) Whole-mount *in situ* hybridization of *Wnt11* expression in E11.5 wildtype (K) and *Gas1* mutant (L,M) kidneys. Red arrowhead (Inset, M) indicates ectopic mesonephric *Wnt11* expression. (N-P) FISH for *Etv5* (yellow) coupled with RET antibody detection (cyan) in E11.5 metanephros coronal sections from wildtype (N) and *Gas1^-/-^*(O,P) embryos. (Q) Quantitation of *Etv5* UB fluorescence intensity in E11.5 wildtype (n=11) and *Gas1^-/-^* (n=9) metanephros sections (3 sections/kidney) normalized to ureteric bud (UB) area. (R- T) FISH for *Wnt11* (yellow) coupled with RET antibody detection (cyan) in E11.5 metanephros coronal sections from wildtype (R) and *Gas1^-/-^*(S,T) embryos. (U) Quantitation of *Wnt11* UB fluorescence signal intensity in E11.5 wildtype (n=13) and *Gas1^-/-^* (n=10) metanephros sections (3 sections/kidney) normalized to ureteric bud (UB) area. Scale bars (C,N) 50μm; (H) 100μm. P- values were determined by a two-tailed Student’s t-test: non-significant: p>0.05 and significant: p≤0.05.

*Gas1* can modulate RET signaling in multiple contexts (Biau et al., 2013; Cabrera et al., 2006; Li et al., 2019; Lopez-Ramirez et al., 2008). To determine if *Gas1* deletion alters downstream RET activation, we collected wildtype and *Gas1^-/-^* embryos at the onset of branching morphogenesis and performed WISH for *Etv5* (Fig.5H-J) and *Wnt11* (Fig.5K-M), two transcriptional targets of RET signaling (Lu et al., 2009; Majumdar et al., 2003). In wildtype embryos, *Etv5* is expressed in the UB epithelium and surrounding MM (Fig. 5H). In *Gas1^-/-^* embryos we observe two categories of *Etv5* expression: 1) a mild reduction in domain of metanephric *Etv5* expression (Fig. 5I), and 2) an apparent loss of *Etv5* expression in the UB and MM (Fig. 5J). Similarly, *Wnt11* is expressed in the UB in E11.5 wildtype embryos (Fig. 5K), with either a mild reduction in the *Wnt11* expression domain (Fig. 5L) or a loss of *Wnt11* expression in *Gas1^-/-^* metanephros (Fig. 5M). To quantify changes in *Wnt11* and *Etv5* expression in the metanephros, we performed section FISH coupled with RET antibody detection in E11.5 wildtype and *Gas1^-/-^* coronal sections (Fig. 5N-U). Consistent with our WISH data, *Gas1^-/-^* metanephroi display variable reductions in *Etv5* expression in both the epithelium and the mesenchyme as well as variably reduced *Wnt11* expression in the UB (Fig. 5Q,U; Fig. S6Q), suggesting that the spectrum of renal agenesis phenotypes observed upon *Gas1* loss results from variable reductions in RET signaling. Interestingly, we identified ectopic *Etv5* and *Wnt11* expression in *Gas1^-/-^* mesonephroi (Fig. 5I,J arrowheads; Fig. 5M, inset), suggesting increased RET signaling in an adjacent tissue.

To investigate the mechanisms underlying GAS1 modulation of RET signaling in metanephros branching morphogenesis, we collected E12.0 wildtype and *Gas1^-/-^* embryos and performed antibody detection for GAS1, co-stained with RET and HSPG2 (Perlecan) to identify the UB epithelium and basement membrane, respectively (Fig. 6A-J). While *Gas1* (B-GAL+) is exclusively expressed in the MM in *Gas1^lacZ/+^* reporter mice, we identified broad GAS1 expression that localizes to the MM and UB epithelium (RET+) in wildtype embryos (Fig. 6A- E). Importantly, GAS1 expression is absent in *Gas1^-/-^* metanephros sections, confirming antibody specificity (Fig. 6G,J). Epithelial GAS1 expression localizes to the cell membrane, with the most robust signal at the basal surface adjacent to the basement membrane (HSPG2+) and luminal side of the UB, similar to the regions of RET expression. Given this epithelial GAS distribution, we hypothesized that GAS1 can be secreted from the MM and accumulate in the UB. To test this, we cultured primary MM cells from E18.5 wildtype and *Gas1^-/-^* metanephros and performed GAS1 immunoprecipitation from the cell supernatants (Fig. 6K). We identified a band at the correct size for GAS1 in wildtype MM supernatants, which was absent in *Gas1^-/-^*MM supernatants, suggesting that GAS1 can be released from the MM. Previous studies identified physical interactions between GAS1 and RET (Cabrera et al., 2006; Li et al., 2019), therefore, we performed fluorescent *in situ* proximity ligation assay analysis coupled with CDH1 antibody detection in E11.5 wildtype and *Gas1^-/-^* metanephros coronal sections to investigate potential interactions of RET and GAS1 in the ureteric bud (Fig. 6L-Y). To validate the assay, we performed PLA for RET and GFRA1, where physical interactions have been described previously (Jing et al., 1996; Treanor et al., 1996). We could detect GFRA1-RET PLA+ puncta in the UB epithelium in wildtype metanephros sections (Fig. 6L-N). In *Gas1* mutants kidneys, we identified similar levels of RET-GFRA1 PLA+ puncta in the UB compared to wildtype embryos, indicating that *Gas1* deletion does not significantly alter RET-GFRA1 interactions (Fig. 6O-Q,R). Next, we performed PLA for potential RET and GAS1 physical interactions in wildtype and *Gas1* mutant kidneys (Fig. 6S-Y). Importantly, few PLA puncta were detected in *Gas1^-/-^* UBs, whereas the levels of RET-GAS1 PLA signal in wildtype embryos was significantly increased (Fig. 6Y). These data suggest a mechanism by which MM-secreted GAS1 can form physical interactions with RET in the UB epithelium to regulate early renal branching morphogenesis.

**Figure 6.**
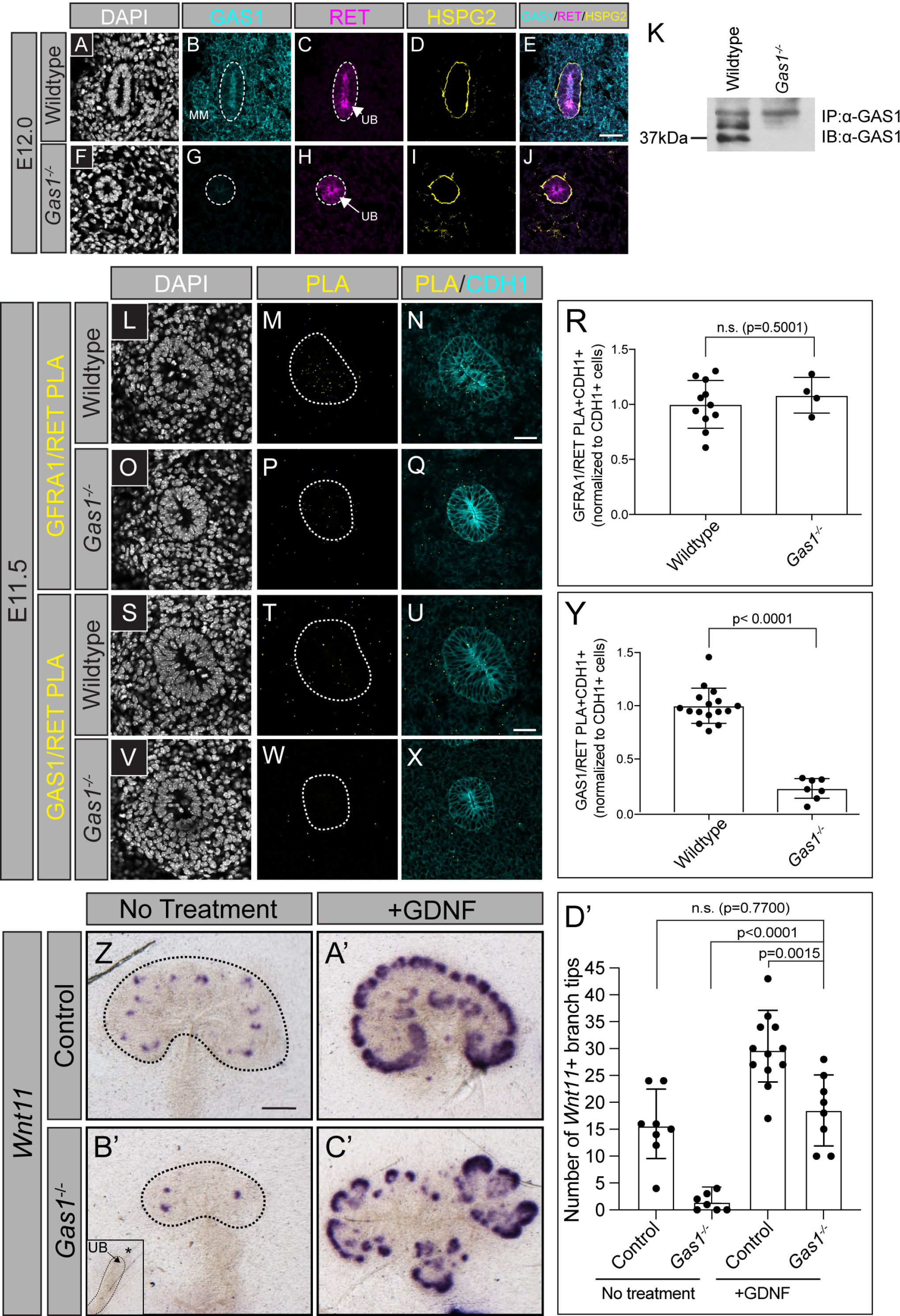
GAS1 is secreted from the MM and accumulates in the UB epithelium. (A-J) Immunofluorescent detection of DAPI (gray; A,F), GAS1 (cyan; B,G), RET (magenta; C,H), HSPG2 (yellow; D,I) in metanephros coronal sections from E12.0 wildtype (A-E) and *Gas1^-/-^*(F-J) embryos. GAS1/RET/HSPG2 merged images are shown (E,J). Dotted lines circle the ureteric bud (UB; B,G,C,H). (K) Western blot detection of immunoprecipitated endogenous GAS1 from supernatants collected from cultured wildtype and *Gas1* mutant metanephric mesenchyme (MM) cells from E18.5 kidneys (n=3 technical replicates). (L-Q) Representative images of Proximity Ligation Assay (PLA) detection of GFRA1/RET PLA+ signal (yellow; M,P) merged with CDH1 (cyan; N,Q) antibody detection in metanephros coronal sections from E11.5 wildtype (L-N) and *Gas1^-/-^* (O-Q) embryos. DAPI labels nuclei (gray; L,O). Dotted lines mark the UB (M,P). (R) Quantitation of the total number of GFRA1/RET PLA+ puncta (normalized to CDH1+ cells; 3 sections/kidney) in E11.5 wildtype (n=11) and *Gas1^-/-^* (n=4) metanephros. (S-X) Representative images of PLA detection of GAS1/RET PLA+ signal (yellow; T,W) merged with CDH1 (cyan; U,X) antibody detection in metanephros coronal sections from E11.5 wildtype (S-U) and *Gas1^-/-^* (V-X) embryos. DAPI labels nuclei (gray; S,V). Dotted lines mark the UB (T,W). (Y) Quantitation of the total number of GAS1/RET PLA+ puncta (normalized to CDH1+ cells; 3 sections/kidney) in E11.5 wildtype (n=16) and *Gas1^-/-^* (n=7) metanephros. (Z-C’) Whole-mount *in situ* hybridization of *Wnt11* expression in wildtype (Z,A’) and *Gas1^-/-^* (B,C’) kidney explants cultured for 4 days in the presence or absence of recombinant murine GDNF (75ng/mL). Dotted lines denote the metanephros (Z’B’) or UB (B’; inset). (D’) Quantitation of the total number of *Wnt11*-expressing UB branch tips in non-treated and GDNF-treated control (wildtype or *Gas1^+/-^*; n=8 non-treated, n=12 GDNF-treated) and *Gas1^-/-^* (n=7 non-treated, n=8 treated) kidney explant cultures. Scale bar (F,N) 25μm; (Z) 250μm. P-values were determined by a two-tailed Student’s t-test (R,Y) or an ordinary one-way ANOVA with Tukey’s multiple comparisons test (D’): p>0.05 and significant: p≤0.05.

To test whether RET pathway activation can rescue the branching morphogenesis defects observed in *Gas1* mutant kidneys, we cultured E11.5 control (wildtype or *Gas1^+/-^*) and *Gas1^-/-^*kidney explants for 4 days in the presence or absence of GDNF ligand. To confirm effective GDNF stimulation, we performed WISH for *Wnt11* on day 4 kidney explants to readout downstream RET pathway activation (Fig. 6Z-D’). In control untreated kidneys, we identified many *Wnt11-*expressing UB tips (Fig. 6Z,D’), whereas untreated *Gas1^-/-^* kidney explants had very few *Wnt11+* branches or completely lacked UB branching (Fig. 6B’,D’), indicative of hypoplasia and agenesis, respectively. Addition of exogenous GDNF results in increased *Wnt11* expression in control explants, which showed a 1.89-fold increase in the number of *Wnt11+* branch tips compared with non-treated control explants (Fig. 6A’,D’). Notably, GDNF-mediated RET activation was able to induce UB branching in 100% of *Gas1^-/-^* kidneys, which exhibited a 12.94-fold increase in the number of *Wnt11+* branches compared with non-treated *Gas1^-/-^*metanephros and branched to a similar degree of non-treated control kidneys (Fig. 6C’,D’). In contrast, *Gas1^-/-^* kidney explants treated with Smoothened agonist (SAG), which stimulates HH signaling downstream of GAS1, did not display a significant increase in UB branching, despite induction of HH pathway activity as demonstrated by WISH for *Gli1* expression (Fig. S7A-E). These data indicate that the UB branching defects and decreased RET signaling observed following *Gas1* deletion can be selectively rescued by activation of the RET pathway.

## Discussion

In this study we identified HH-dependent and HH-independent contributions of GAS1 to early mammalian kidney morphogenesis (Figure 7). We found that *Gas1* is expressed in the early kidney mesenchyme and is selectively required for proper mesonephro- and metanephrogenesis in a genetic background-dependent manner. Specifically, *Gas1* deletion results in ectopic mesonephric tubule formation, a phenotype that correlates with decreased HH signaling in the mesonephros. Further, *Gas1* deletion results in a transient decrease in metanephric mesenchyme cell proliferation and reduced RET target gene expression, which contributes to early renal hypoplasia and agenesis. Mechanistically, our data indicate that secreted GAS1 binds RET in the ureteric bud epithelium to promote RET pathway activation and subsequent UB branching morphogenesis. Importantly, the metanephros defects observed in *Gas1* mutants can be rescued by increasing RET pathway activity via exogenous GDNF stimulation in *Gas1* mutant explant cultures. Overall, these data highlight essential, multifunctional roles for GAS1 in early kidney morphogenesis.

**Figure 7.**
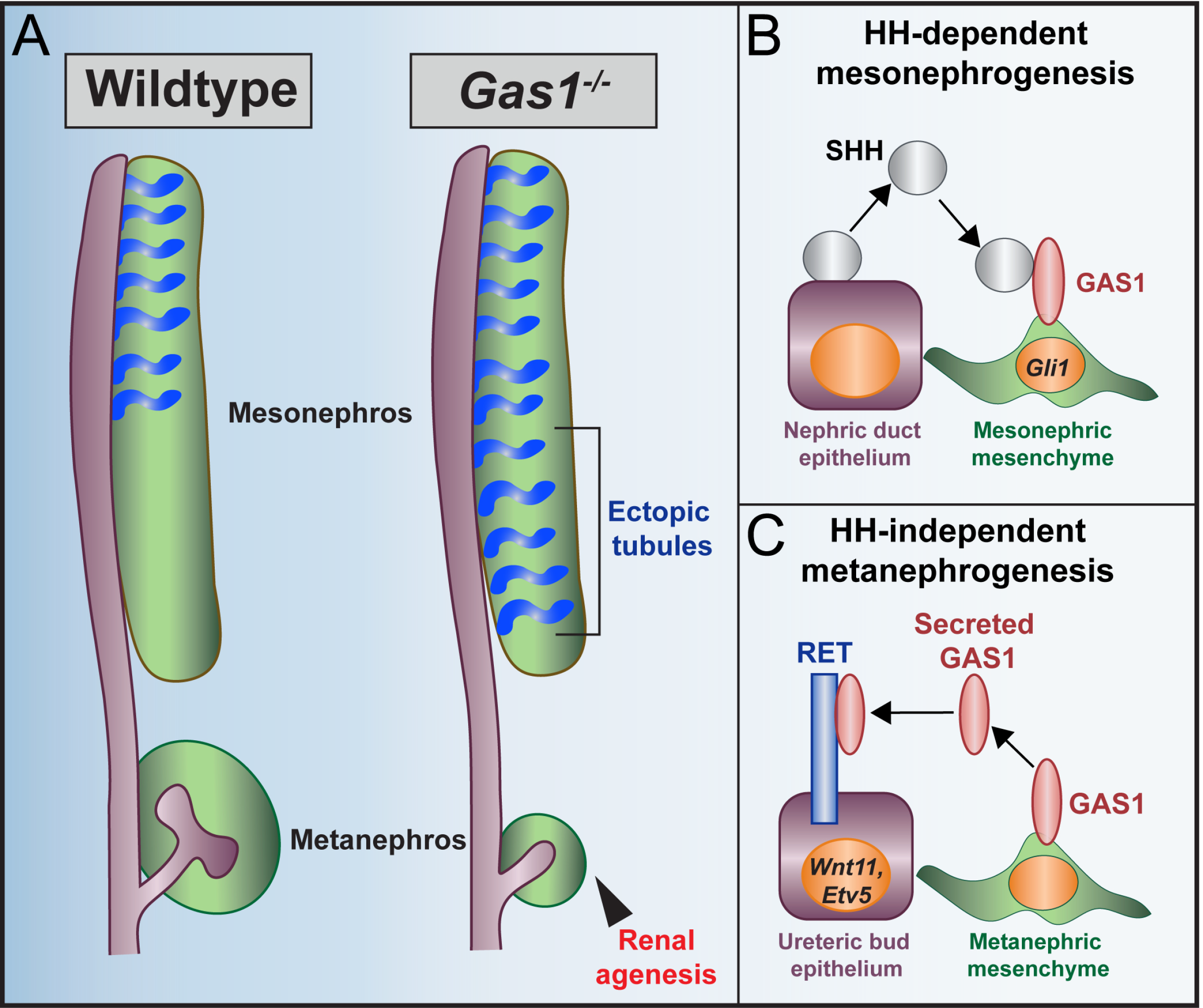
GAS1 is a multifunctional regulator of early kidney morphogenesis. (A) Summary of GAS1 contributions in mesonephrogenesis and metanephrogenesis. *Gas1* deletion results in ectopic caudal mesonephric tubules and variable renal agenesis. (B) Proposed mechanism for GAS1 regulation of mesonephros development: SHH ligand (gray) is secreted from the nephric duct epithelium (purple) and interacts with GAS1 (red) in the adjacent mesonephric mesenchyme (green) to drive HH pathway activation (*Gli1* expression). (C) Proposed mechanism for GAS1 regulation of metanephros development: GAS1 (red) is secreted from the metanephric mesenchyme (green) and interacts with RET receptors (blue) in the ureteric bud epithelium (purple) to promote RET pathway activation (*Wnt11* and *Etv5* expression).

### Unique role for GAS1 in HH-dependent mesonephrogenesis

Despite the expression of all three HH co-receptors in developing mouse kidneys, we find here that *Gas1*, but not *Cdon* and *Boc*, is essential for HH-dependent mesonephrogenesis.

Notably, there are precedents for distinct contributions of these co-receptors to the development of other tissues. For example, during craniofacial development, the co-receptor BOC plays a partially antagonistic role that is functionally distinct from GAS1 and CDON (Echevarria-Andino and Allen, 2020). CDON also restricts HH signaling during optic vesicle patterning in zebrafish (Cardozo et al., 2014). Further, during limb development, only the loss of *Gas1*, but not *Cdon* and *Boc*, results in digit specification defects (Allen et al., 2011).

One explanation for these different functional contributions to kidney development is the structural differences between GAS1, a GPI-anchored protein resembling GDNF receptors (Cabrera et al., 2006), and CDON and BOC, single pass transmembrane proteins that are members of the Ig superfamily (Kang et al., 1997; Kang et al., 2002). It is also possible that the level of expression and cell type-specific expression of each HH co-receptor affects their relative contribution (e.g., mesenchymal *Cdon* expression appears more restricted than either *Gas1* or *Boc*, while *Cdon* and *Boc,* but not *Gas1,* are expressed in both the kidney epithelium and metanephric mesenchyme).

Evaluation of the combined contributions of all three HH co-receptors in later embryogenesis has been hindered by the early embryonic lethality of *Gas1;Cdon:Boc* germline mutants (Allen et al., 2011). However, the successful development of conditional alleles for both *Cdon* and *Gas1* (Bae et al., 2020; Jin et al., 2015) provides an opportunity to assess their combined contribution in the kidney mesenchyme. These will be particularly informative experiments since *Gas1* deletion does not completely abrogate HH pathway activity during mesonephros development, and since a role for *Cdon* in HH-dependent digit specification was only revealed in a *Gas1;Boc* null background (Echevarria-Andino et al., 2022).

### GAS1 regulates multiple signaling pathways during kidney development

GAS1 functions in several key developmental signaling pathways, including HH, RET, FGF and NOTCH (Allen et al., 2011; Allen et al., 2007; Biau et al., 2013; Cabrera et al., 2006; Izzi et al., 2011; Kann et al., 2015; Li et al., 2019; Lopez-Ramirez et al., 2008; Marczenke et al., 2021; Martinelli and Fan, 2007a; Seppala et al., 2007). GAS1-mediated regulation of these pathways is thought to occur via direct physical interactions between GAS1 and different signaling molecules, including secreted HH ligands and various membrane receptors such as PTCH1, RET, and NOTCH1 (Cabrera et al., 2006; Izzi et al., 2011; Lee et al., 2001; Li et al., 2019; Marczenke et al., 2021). While GAS1 binds HH ligands with high affinity (Huang et al., 2022; Lee et al., 2001), lower affinity interactions have been reported between GAS1 and RET receptors (Cabrera et al., 2006; Li et al., 2019; Rosti et al., 2015). In regions with overlapping GAS1/RET/HH expression domains, higher affinity interactions between GAS1 and HH might drive HH pathway activation, while restricting RET signaling. Notably, *Shh* and *Ret* show similar expression patterns in the kidney epithelium prior to UB induction but become more restricted to the mesonephros/distal ureter (*Shh*) and UB tips (*Ret*) around E11.5, while *Gas1* maintains broad mesenchymal expression throughout these tissues. Consistent with this notion, we find that *Gas1* deletion results in decreased HH signaling and ectopic expression of *Ret* target genes in early mesonephrogenesis.

Our data indicate that GAS1 can be secreted in the metanephros, therefore the requirements for GAS1 in a specific pathway (e.g., HH versus RET) could depend on whether it is acting in a cell-autonomous manner (i.e., membrane-tethered) or in a non-cell autonomous fashion (i.e., secreted). GAS1 may also act on multiple signaling pathways simultaneously in a single tissue during development, as suggested for gut development, forebrain development and dentition patterning (Biau et al., 2013; Marczenke et al., 2021; Seppala et al., 2022).

Alternatively, GAS1 may function in still other pathways. For example, *Foxc1, Slit2,* and *Robo2* mutant mesonephroi resemble the phenotypes we observe in *Gas1^-/-^* mesonephros (Grieshammer et al., 2004; Kume et al., 2000), suggesting that GAS1 may also contribute to these pathways during mesonephrogenesis. Further studies dissecting the contributions of GAS1 to multiple pathways will also need to consider whether GAS1 is working independently in these pathways, or perhaps acting to integrate multiple signals received by a cell. Here, a better understanding is needed of the subcellular locations of GAS1 protein with respect to its interacting partners, as well as potential competition for GAS1 binding with these proteins, and the precise timing and duration of the signaling amongst these pathways.

### GAS1 as a secreted protein

Secreted HH-binding proteins are essential for proper HH pathway activity. SCUBE2 promotes the release and spread of HH ligands through tissues (Creanga et al., 2012; Tukachinsky et al., 2012), while HHIP, a secreted HH pathway antagonist, acts non-cell autonomously to restrict HH signaling in the developing ventral neural tube and lung (Holtz et al., 2015). There is also evidence that the HH-coreceptor BOC can be released from the cell surface, although membrane tethering of BOC and CDON is essential for HH-dependent neural patterning in the developing chicken neural tube (Song et al., 2015). Here, we identified GAS1 secretion from the kidney mesenchyme during development, however potential functional roles for secreted GAS1 have not been explored.

Previous studies described roles for soluble GPI-anchored proteins (GPI-AP) in multiple developmental processes. For example, release of the GPI-AP CRIPTO is required for non-cell-autonomous activation of Nodal signaling and plays roles in axial midline development (Chu et al., 2005; Lee et al., 2016; Parisi et al., 2003; Watanabe et al., 2007; Yan et al., 2002). Further, glycerophosphodiester phosphodiesterase 2 (GDE2)-dependent cleavage and inactivation of the GPI-AP, RECK regulates motor neuron differentiation through Notch inhibition (Muraguchi et al., 2007; Park et al., 2013; Sabharwal et al., 2011). Interestingly, two additional GDE proteins, GDE6 and GDE3, are expressed during development and regulate multiple aspects of neurogenesis (Dobrowolski et al., 2020; McKean et al., 2023), a tissue where GAS1 has well-described roles in promoting HH-dependent neural patterning. Further reports show that ADAM10 and 17 enzymes can release GAS1 in cultured mesangial cells (van Roeyen et al., 2013). Whether GDE, ADAM, or additional cleavage proteins mediate GAS1 release in the developing kidney remains to be investigated.

This study also raises questions about GAS1 presentation to various binding partners (e.g., secreted or membrane-tethered), including how its presented to neighboring cells (e.g., *cis* versus *trans*). Membrane-bound and soluble GPI-anchored GDNF receptors can activate RET through *cis* and *trans* interactions, respectively, (Fleming et al., 2015; Jing et al., 1996; Ledda et al., 2002; Paratcha et al., 2001; Yu et al., 1998); however, *trans Gfra1* signaling is dispensable for normal development in mice (Enomoto et al., 2004). Similarly, GAS1 requires membrane tethering to promote delta-like 1 (DLL-1)-mediated activation of NOTCH receptors (Marczenke et al., 2021). The development of genetic tools to selectively express membrane-anchored or secreted GAS1 will be essential to determine the relative contributions of these distinct GAS1 variants to kidney development and to a host of key developmental signaling pathways.

## Acknowledgements

We thank members of the Allen Lab for their feedback and support with this project. We also acknowledge Dr. Cristina Cebrian for technical assistance with the kidney explant culture experiments used in this study. We also thank Dr. Greg Dressler for providing valuable suggestions and reagents. Finally, we acknowledge the Biomedical Research Core Facilities Microscopy Core for providing access to confocal microscopy equipment.

## Author Contributions

Conceptualization, N.E.F., and B.L.A.; Methodology, N.E.F., and B.L.A.; Investigation, N.E.F.; Formal Analysis, N.E.F.; Writing – Original Draft, N.E.F.; Writing – Review & Editing, N.E.F. and B.L.A.; Funding Acquisition, N.E.F. and B.L.A.; Resources, B.L.A.; Supervision, B.L.A.

## EXPERIMENTAL MODEL AND SUBJECT DETAILS

### Animals

*Gas1*^lacZ^ (Martinelli and Fan, 2007a), *Cdon^lacZ-2^*(Cole and Krauss, 2003), *Cdon^lacZ-1^* (Cole and Krauss, 2003), *Boc*^AP^ (Zhang et al., 2011), *Gli1*^lacZ^ (Bai and Joyner, 2001) and *Ret* (Schuchardt et al., 1994) mice have been described previously. *Gas1* mice were maintained on congenic C57BL6/J and 129S4/SvJaeJ genetic backgrounds. *Cdon* and *Boc* mice were maintained on a congenic C57BL6/J background. *Gli1*, *Ret* and *Cdon^lacZ-1^* mice used for expression analysis were maintained on a mixed 129S4/SvJaeJ/C57BL/6J background. Noon on the day of the vaginal plugs was considered E0.5. All experiments were conducted using both male and female embryos. All animal procedures were reviewed and approved by the Institutional Animal Care and Use Committee (IACUC) at the University of Michigan.

## METHOD DETAILS

### Section Immunofluorescence

Immunofluorescence was performed as described previously (Allen et al., 2011). Kidneys were harvested at various developmental stages and fixed in 4% paraformaldehyde for 20 min on ice, washed 3 x 5 min in 1X PBS (pH 7.4) and cryoprotected in 1X PBS + 30% sucrose for 24 h.

Tissues were embedded in OCT and cryo-sectioned on a Leica CM1850 cryostat at a thickness of 12 μm. Sections were washed with 1X PBS and blocked for 1 h at room temperature in blocking buffer (1X PBS, 10% donkey serum, 0.3% Triton-X for goat anti-GAS1 and goat anti- GFRa1 antibodies; all other antibodies were blocked in 1X PBS, 3% bovine serum albumin, 1% heat inactivated sheep serum, and 0.1% Triton-X 100). Slides were incubated in the indicated primary antibodies diluted in buffer overnight at 4°C in a humidified chamber. All AlexaFluor secondary antibodies were diluted 1:500 in blocking buffer and applied to the sections for 1 h at room temperature. To label nuclei, DAPI was applied for 10 min at a concentration of 1:30,000 in blocking buffer. Sections were washed 3 x 5 min in 1X PBS and mounted using Immu-mount mounting medium. All images were obtained using a Leica SP5 upright confocal.

### Whole-mount immunofluorescence

Staining of whole kidneys was performed as described previously (Barak and Boyle, 2011). Kidneys were dissected in 1X PBS and fixed for 20 minutes in 4% PFA. They were washed 3 x 5 min in 1X PBS and blocked for 1 hour in blocking solution (1X PBS, 0.1% Triton-X100, 10% sheep serum). Kidneys were incubated in E-Cadherin (1:100; BD Biosciences Cat#610181) antibody diluted in block solution overnight at 4°C. The following day, kidneys were washed 4 x 1 h in 1X PBS + 0.1% Triton-X 100. AlexaFluor secondary antibodies were applied to tissues at a concentration of 1:500 and incubated overnight at 4°C. Kidneys were washed as described previously and images were collected using a Nikon SMZ1500 stereomicroscope.

### Generation of Riboprobes

RNA was extracted from E15.5 wildtype mouse kidneys using the Purelink RNA mini-kit (Invitrogen, Cat#12183025) and reversed transcribed using the High-Capacity cDNA Reverse Transcriptase Kit (Applied Biosystems, Cat# 4368814). Gene specific primers containing a T7 promoter sequence were designed for *Wnt11* (Forward: CCGCCACCATCAGTCACACCAT; Reverse: TAATACGACTCACTATAGGGGACGTAGCGCTCCACCGTGC), *Gfra1* (Forward: CCTCGATGCAGCCAAGGCC; Reverse: TAATACGACTCACTATAGGGTGGGAATCTCATTCTCAGAG) and *Etv5* (Forward: AGCCCACCATGTATCGAGAG; Reverse: TAATACGACTCACTATAGGGTCAAAGGGCAAGCTTTAGGA) using Primer3 plus. cDNA was amplified by PCR and subcloned into TOPO 2.1 vector and linearized by restriction enzyme digestion. Digoxigenin labeled antisense riboprobes were synthesized by *in vitro* transcription using a DIG RNA labeling kit (Roche, Cat#11277073910), followed by purification using Micro Bio-Spin P-30 gel columns (Bio-Rad, Cat#7326223). The purified RNA was quantified using a Nanodrop and diluted to a concentration of 20 ng/μl in hybridization buffer (50% formamide, 5X SSC (pH 4.5), 50 μg/ml yeast tRNA, 1% SDS, 50 μg/ml heparin).

### Whole mount *in situ* hybridization

*In situ* hybridization was performed as described previously (Allen et al., 2011), (Wilkinson, 1992). Briefly, kidneys were harvested and fixed in 4% paraformaldehyde for 24 h at 4°C. Tissues were dehydrated through a 1X PBST + methanol series (25% methanol, 50% methanol and 75% methanol) and stored at -20°C in 100% methanol for up to 3 months. Embryos were washed in 1X PBS + 0.1% Tween, bleached for 1 h in 3% hydrogen peroxide, and digested with 10 μg/mL Proteinase-K for 5 min. Tissues were hybridized with the indicated DIG-labeled riboprobes diluted to 1 ng/ml in hybridization buffer (50% formamide, 5X SSC, pH 4.5, 50 μg/ml yeast tRNA, 1% SDS, 50 μg/ml heparin) at 70°C for 16-20 h. Kidneys were incubated in alkaline phosphatase-conjugated anti-DIG antibody (1:4000, Roche, Cat#11093274910). For *in situ* detection, kidneys were incubated in BM purple substrate for up to 48 h depending on the probe. Tissues were post-fixed in 4% PFA + 0.1% glutaraldehyde, cleared in 1X PBST + 80% glycerol and imaged using a Nikon SMZ1500 stereomicroscope. *Etv5, Wnt11,* and *Gfra1* RNA *in situ* probes were designed from E15.5 mouse kidney cDNA as indicated above. *Pax2, Gli1,* and *Ret* DIG-labelled RNA probes were gifted from Andrew McMahon (McMahon et al., 2008)

### Section *in situ* hybridization

Kidneys were harvested and fixed in 4% paraformaldehyde for 24 h at 4°C. The next day, tissues were processed according to the immunofluorescence protocol and sectioned at a thickness of 20 μm. Sections were thawed and post-fixed in 4% PFA for 10 min, followed by 3 x 5 min washes in 1X PBS, and digested for 2 minutes in 10 μg/ml of proteinase K. Sections were fixed again in 4% PFA for 5 min and incubated in acetylation buffer (Water, triethanolamine, hydrochloric acid, and acetic anhydride) for 10 min. Sections were washed and dehydrated through a series of solutions (1X PBS, 0.85% NaCl, 70% ethanol, 95% ethanol). *Gli1* probe was diluted to 1 ng/μl in hybridization buffer (50% formamide, 5X SSC, pH 4.5, 50 μg/ml yeast tRNA, 1% SDS, 50 μg/ml heparin) and slides were incubated at 70°C for 16-20 h. The next day, sections were washed with 1X SSC (saline sodium citrate, pH4.5) + 50% formamide at 65°C for 30 min, followed by TNE buffer (10mM Tris pH7.5, 500mM NaCl, 1mM EDTA) wash for 10 min at 37°C, then incubated in 5 μg/ml RNase A in TNE at 37°C for 15 min. Slides were washed in a series of buffers (TNE, 2XSSC, 0.2XSSC, MBST), blocked in MBST + 10% HISS + 2% BMB for 2 h at room temperature and incubated in alkaline phosphatase-conjugated anti-DIG antibody (1:4000, Roche, Cat#11093274910) diluted in blocking buffer until signal development. Slides were mounted with Glycergel mounting medium and imaged using a Nikon-E800 microscope.

### X-gal staining

Embryos were collected in 1X PBS (pH 7.4) and fixed (1% formaldehyde, 0.2% glutaraldehyde, 2mM MgCl2, 5mM EGTA, 0.02% NP-40) for 1 h at room temperature. Tissues were cryoprotected in 1X PBS + 30% sucrose overnight at 4°C, embedded in OCT, and sectioned using a Leica CM1850 cryostat at a thickness of 20 μm. Sections were incubated in staining solution (5mM K3Fe(CN)6, 5mM K4Fe(CN)6, 2mM MgCl2, 0.01% Na deoxycholate, 0.02% NP- 40, 1mg/mL X-gal) until signal developed. Slides were washed 3 x 5 min in 1X PBS and counterstained with nuclear fast red for 5 min. Sections were washed in 1X PBS, dehydrated through a series of ethanol and xylenes (70% ethanol, 100% ethanol, 100% xylenes) and mounted using Permount mounting medium (Fisher Chemical, Cat#SP15100). Sections were imaged using a Nikon E-800 microscope.

### Alkaline phosphatase staining

Wildtype and *Boc^AP^* embryos were dissected in 1X PBS and fixed for 1 h in 4% PFA. Embryos were cryoprotected in 1X PBS + 30% sucrose for 24 h, embedded in OCT, and sectioned on a Leica cryostat at 20 μm. Endogenous alkaline phosphatase was deactivated by incubating slides in 1X PBS for 30 min at 70°C. BM purple substrate was applied to the sections and incubated for 3-4 h at 37°C. Following staining, slides were processed as described for X-gal staining and imaged on a Nikon E-800 microscope.

### Fluorescent RNAscope *in situ* hybridization

Embryos were dissected in 1X PBS and fixed in 10% neutral buffered formalin (NBF) for 20-24 h at room temperature. The following day, tissues were cryoprotected in 30% sucrose for 24 h, embedded in OCT, and sectioned at a thickness of 10 μm. Fluorescent RNAscope (ACD; Multiplex Fluorescent Assay Kit V2, Cat#323110) was performed according to the manufacturer’s instructions. Antigen retrieval was performed for 15 min at 100°C in a steamer and sections were treated with protease plus for 1 min, followed by incubation with the indicated probes. Antibody staining was performed as described previously for section immunofluorescence and slides were imaged on a Leica SP5 upright confocal.

For quantitation, images were manually thresholded using ImageJ and fluorescence signal was measured using the integrated density function normalized to a given area as indicated in the figure legends. Each data point represents one kidney. A minimum of three embryos per genotype and two sections from each kidney were analyzed.

### Kidney explant cultures

Kidney cultures were performed as described previously (Costantini et al., 2011). Briefly, wildtype, *Gas1^+/-^* and *Gas1^-/-^* mutant mouse embryonic kidneys were harvested at E11.5 in cold CO2-independent medium. (Gibco, Cat#18045088). Kidneys were plated on a polyester membrane insert (0.4 μm pore size; 12 mm diameter, Corning, Cat#3460) in a 6-well tissue culture dish. 1.5 mL of kidney culture media (DMEM/F-12 supplemented with 10% calf serum and 1% penicillin-streptomycin glutamine) was applied to the bottom of each well below the membrane. For rescue experiments, recombinant murine GDNF was diluted to 75 ng/ml (Peprotech, cat#4504410UG). Smoothened agonist (SAG, Enzo Life Sciences, Cat#ALX-270- 426-M001) was diluted to a final concentration of 300 nM in kidney culture media. Kidneys were cultured for 4 d at 37°C in a 5% CO2 incubator. Fresh kidney culture media was applied after 2 d of culture. For whole-mount *in situ* hybridization of kidney explant cultures: After 4 d of culture, kidney explant cultures were fixed on the transwell membranes in 4% paraformaldehyde for 24 h at 4°C in a 6-well plate. The next day, kidney explants were washed 3 x 5 min in 1X PBST and processed according to the whole-mount *in situ* hybridization protocol with a 2-min proteinase-K digestion. Kidneys were imaged using a Leica SMZ1500 Stereomicroscope.

### Immunoprecipitation/western blot

Wildtype and *Gas1^-/-^* kidneys were dissected at E18.5 in CO2-independent medium, incubated in 2 mg/ml of collagenase diluted in CO2-independent medium (2 kidneys pooled) for 20 min at 37°C and cells were dissociated by gentle pipetting. Cells were pelleted and resuspended in culture media (DMEM/F12 +10% calf serum + 1% PSG), plated in a 6-well dish and cultured for 2-3 d at 37°C and 5% CO2. Cell supernatants were collected and centrifuged to remove dead cells. For immunoprecipitation, 50 μl of protein-G Dynabeads (Invitrogen,Cat# 10-003-D) were washed in 1X PBS then incubated in goat anti-GAS1 (2 mg, R&D Systems; Cat#AF2644) antibody diluted in 1X PBS for 2 h at room temperature on a rotating platform. Beads were washed twice with 1X PBST, then incubated with 1 mL of cell supernatant for 16-24 h on a rotator at 4°C. The next day, beads were briefly washed 4 times in 1X PBS +0.1% Tween, then 4 times in 1XPBS + 0.1% Triton. Proteins were eluted in 30 μl of 0.1 M citrate (pH 2.0) for 15 min on a hula mixer. To neutralize the pH, 3 μl Tris buffer (pH 8.0) was applied to each sample. For denaturation, samples were boiled at 95°C in 6X Laemmli buffer for 10 min. Proteins were separated by SDS-PAGE on a 10% gel, then transferred onto a Immun-Blot PVDF membrane (Bio-Rad, Cat# 162-0177). Following transfer, membranes were blocked in western blocking buffer (30 g/L bovine serum albumin with 0.2% NaN3 in TBST (Tris-buffered saline, 0.5% tween-20) for 1 h at room temperature. Membranes were incubated in goat anti-GAS1 (1:4000, R&D Systems, AF2644) primary antibody overnight at 4°C. The next day, membranes were washed 4 x 10 min in 1X TBST, followed by a 1-h incubation in peroxidase-conjugated AffiniPure secondary antibodies (1:10000, Jackson ImmunoResearch) and 4 x 10 min washes in 1X TBST. Membranes were incubated in Amersham ECL Prime western blot detection reagent (Cytiva, RPN2232) for 5 min at room temperature and developed using a Konica Minolta SRX- 101A medical film processor.

### Proximity ligation assay

Embryos were dissected in 1X PBS and processed as described above for section immunofluorescent staining. Duolink® Proximity Ligation Assay (PLA; Sigma; DUO92007) was performed according to the manufacturer’s instructions. Sections were blocked (1X PBS+ 10% Donkey serum + 0.3% Triton) for 1 h at 37°C, then incubated with rabbit anti-RET (1:200, Cell Signaling, Cat#3223) and goat anti-GAS1 (1:900, R&D Systems; Cat#AF2544) or goat anti- GFRA1 (1:500, R&D Systems, Cat# AF560) overnight at 4°C. The next day, slides were incubated with PLA Probe Anti-Rabbit PLUS (Sigma; DUO92002) and PLA Probe Anti-Goat Minus (Sigma; DUO92006). After the final washes, slides were processed as described previously for section immunofluorescent staining for detection of mouse IgG2a anti-E-cadherin (1:500, BD Biosciences, Cat# 610181) and imaged on a Leica SP5 upright confocal. For quantitation, PLA puncta were manually counted using the cell counter tool on imageJ and normalized to the total number of E-Cadherin+ cells. A minimum of four kidneys per genotype and three sections/kidney were analyzed.

### Quantitation and statistical analysis

All statistical analyses were performed using GraphPad statistic calculator or GraphPad Prism (www.graphpad.com) and data are reported as mean and standard deviation. Statistical significance was determined using a two-tailed Student’s t*-*test or an ordinary one-way ANOVA with Tukey’s multiple comparisons test. For all experiments, a minimum of three kidneys were analyzed for each genotype. The number of replicates (n) and P-values are listed in each figure. Non-significant: p>0.05 and significant: p≤0.05.

### Kidney area quantitation

Whole embryos and isolated kidneys were imaged using an SMZ1500 Stereomicroscope. Kidney area was measured using the freehand tool in ImageJ and normalized to the embryo length.

Crown-rump length was defined as the top of the crown to the bottom curvature of the embryo at the anterior hindlimb level.

### Immunofluorescence quantitation

The total number of WT1+ kidney mesenchymal cells and PHH3+ cells was manually counted using the cell counter tool feature in ImageJ (Schneider et al., 2012). PHH3+ cells were normalized to total number of E-cadherin+ (Ureteric Bud) or WT1+ (metanephric mesenchyme) cells. For cleaved caspase-3 quantitation, SIX2+ cells were used to identify the metanephric area. In serial sections from the same animal, the number of CC3+ cells per metanephric area was measured using the analyze particle tool in ImageJ. A minimum of three kidneys per genotype and two sections/kidney were analyzed.

---

**Figure S1.**
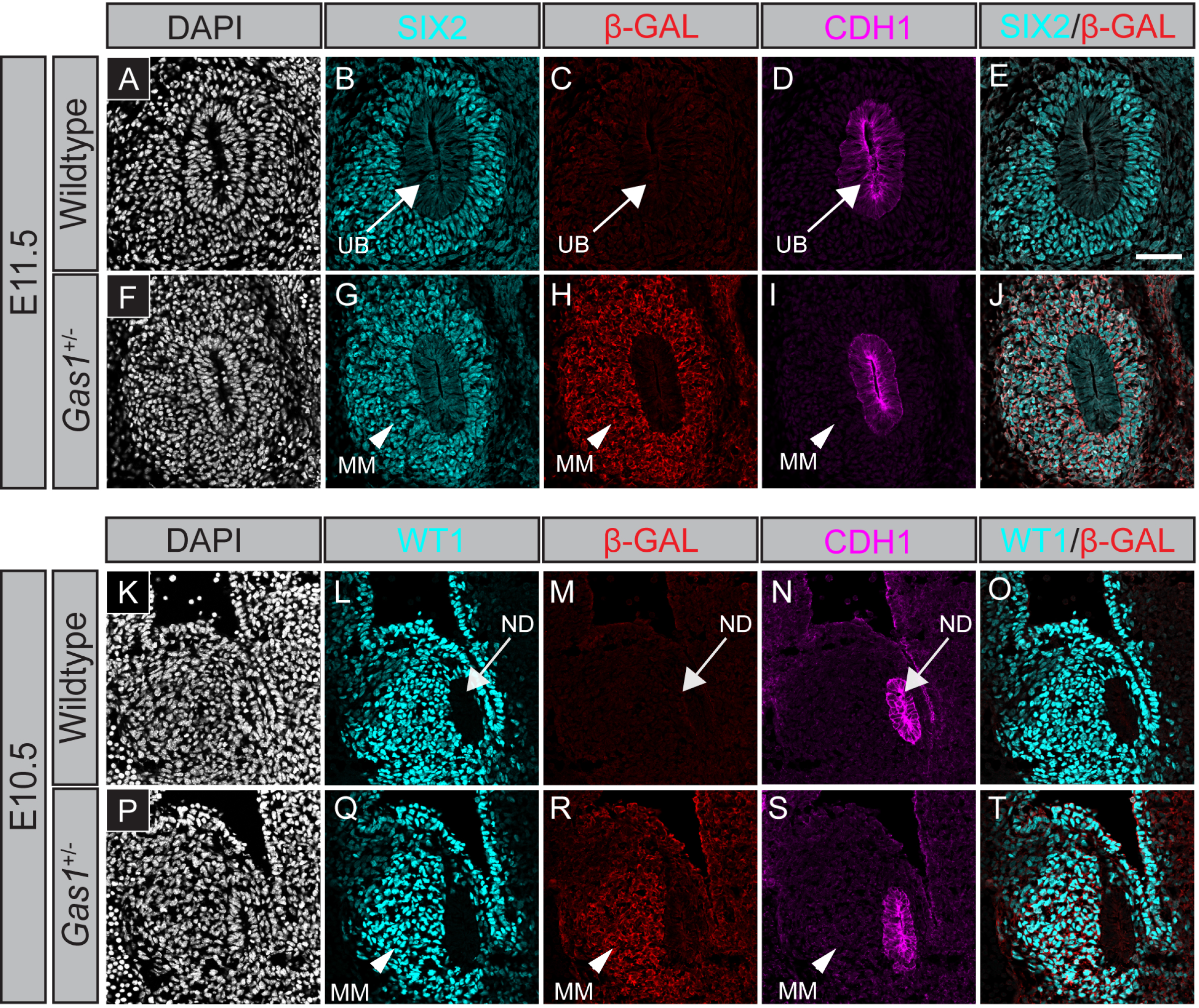
*Gas1* is expressed in the metanephric mesenchyme from the onset of kidney development. (A-T) Immunofluorescent confocal microscope images of coronal metanephros sections from E11.5 (A-J) and E10.5 (K-T) wildtype and *Gas1^lacZ/+^* mouse embryos. Nuclei are labeled with DAPI (A,F,K,P). Antibody detection of SIX2 (cyan; B,G), WT1 (cyan; L,Q), β-galactosidase (β- GAL, red; C,H,M,R) and CDH1 (magenta; D,I,N,S). SIX2/β-GAL (E,J) and WT1/β-GAL (O,T) merged images are shown. Scale bar (E), 50μm. Arrows indicate nephric duct (ND) and ureteric bud (UB). Arrowheads identify metanephric mesenchyme (MM).

**Figure S2.**
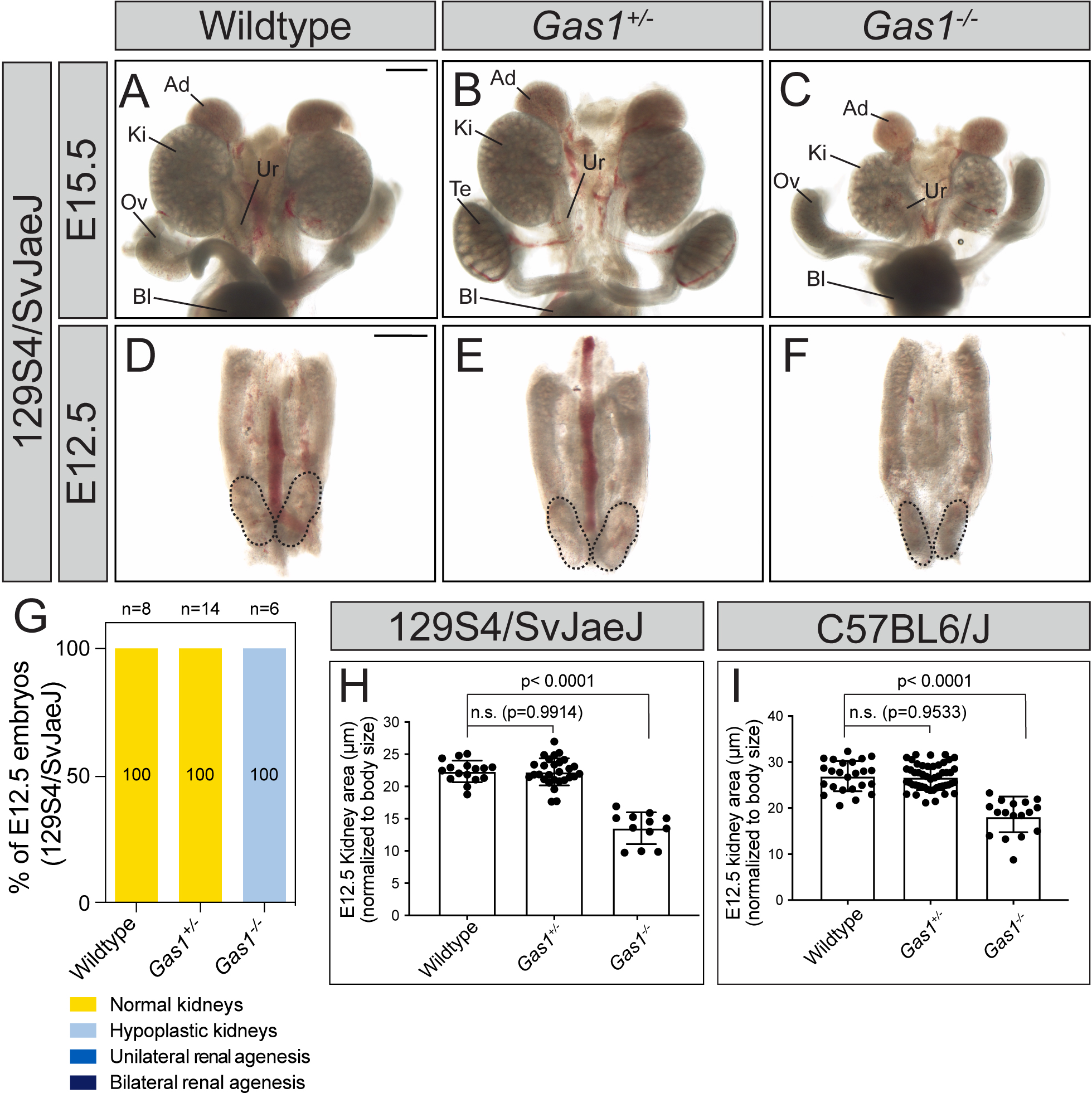
*Gas1* mutant embryos on a congenic 129S4/SvJaeJ background display renal hypoplasia. (A-F) Whole-mount brightfield images of E15.5 (A-C) and E12.5 (D-F) wildtype (A,D), *Gas1^+/-^*(B,E) and *Gas1^-/-^* (C,F) kidneys (marked by dotted lines at E12.5) from mice maintained on a congenic 129S4/SvJaeJ background. (G) Quantitation of the frequency of kidney defects (129S4/SvJaeJ background) in E12.5 (52-56 somites) wildtype (n=8), *Gas1^+/-^* (n=14), and *Gas1^-/-^*(n=6) embryos classified into the following categories: normal kidneys, hypoplastic kidneys, unilateral renal agenesis, or bilateral renal agenesis. (H) Quantitation of kidney area normalized to crown-rump length (in μm) in E12.5 (52-56 somites) wildtype (n=16), *Gas1^+/-^* (n=28) and *Gas1^-/-^*(n=12) embryos from mice on a 129S4/SvJaeJ genetic background. (I) Quantitation of kidney area normalized to crown-rump length (in μm) in E12.5 (52-56 somites) wildtype (n=30), *Gas1^+/-^*(n=43) and *Gas1^-/-^* (n=19) embryos from mice on a C57BL6/J genetic background. Scale bars (A,D) 500μm. P-values were determined by an ordinary one-way ANOVA with Tukey’s multiple comparisons test: p>0.05 and significant: p≤0.05.

**Figure S3.**
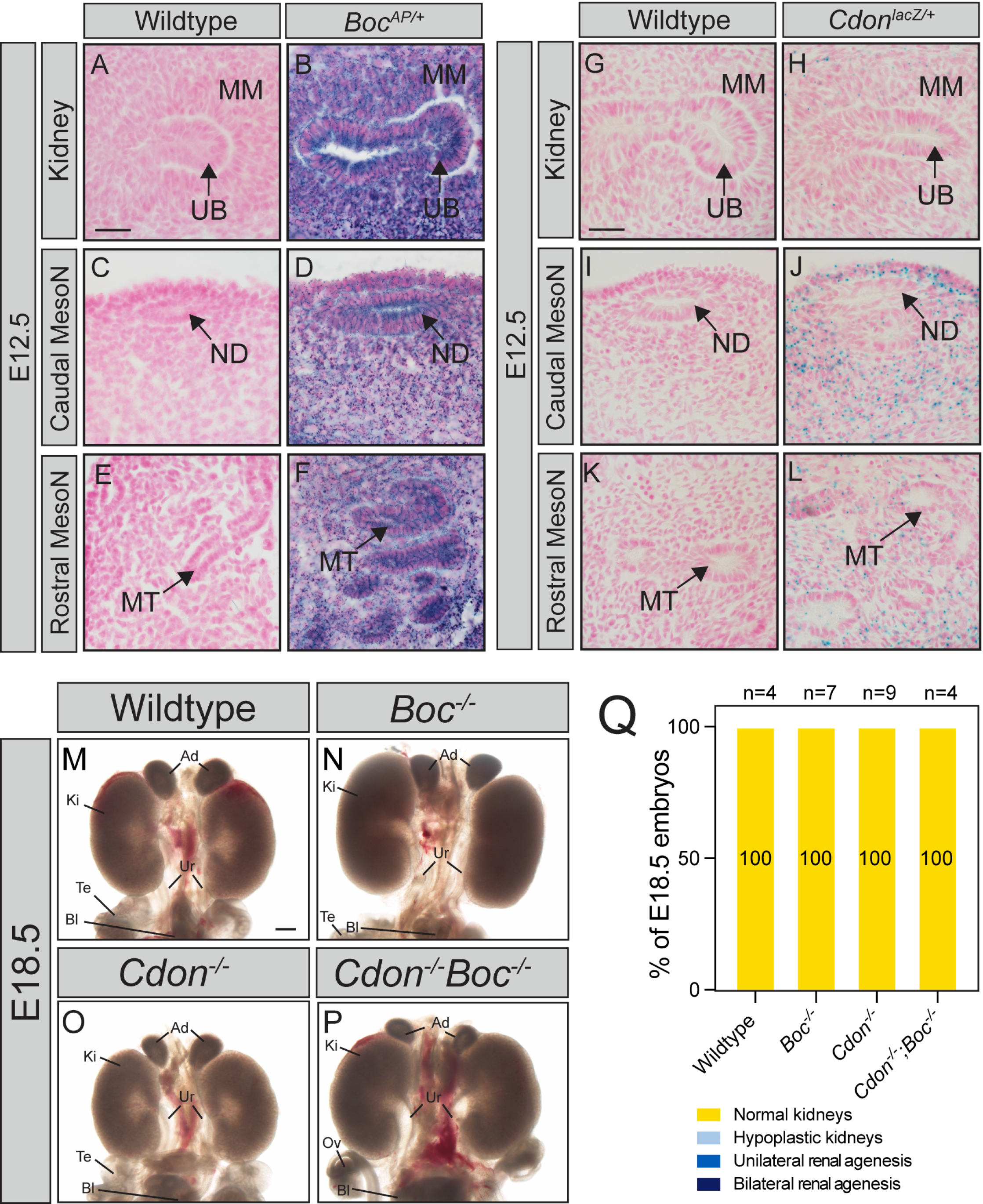
Loss of *Cdon* or *Boc* does not result in renal agenesis. (A-F) Alkaline phosphatase (AP) staining of coronal sections from E12.5 kidney (A,B), caudal mesonephros (C,D) and rostral mesonephros (E,F) in wildtype (A,C,E) and *Boc^AP/+^* (B,D,F) embryos. (G-L) X-gal staining of coronal sections from E12.5 kidney (G,H), caudal mesonephros (I,J) and rostral mesonephros (K,L) in wildtype (G,I,K) and *Cdon^lacZ/+^* (H,J,L) embryos. (M-P) Whole-mount brightfield images of E18.5 wildtype (M), *Boc^-/-^* (N), *Cdon^-/-^*(O) and *Cdon^-/-^;Boc^-/-^* (P) mouse kidneys. (Q) Quantitation of the frequency of renal defects in E18.5 wildtype (n=4), *Boc^-/-^* (n=7), *Cdon^-/-^* (n=9) and *Cdon^-/-^;Boc^-/-^* (n=4) embryos classified into the following categories: normal kidneys, hypoplastic kidneys, unilateral renal agenesis, or bilateral renal agenesis. Scale bars (A,G) 20μm; (M), 500μm. Abbreviations: metanephric mesenchyme (MM), ureteric bud (UB), nephric duct (ND), mesonephric tubules (MT), ureter (Ur), adrenal gland (Ad), kidney (Ki), testis (Te), ovary (Ov), bladder (Bl).

**Figure S4.**
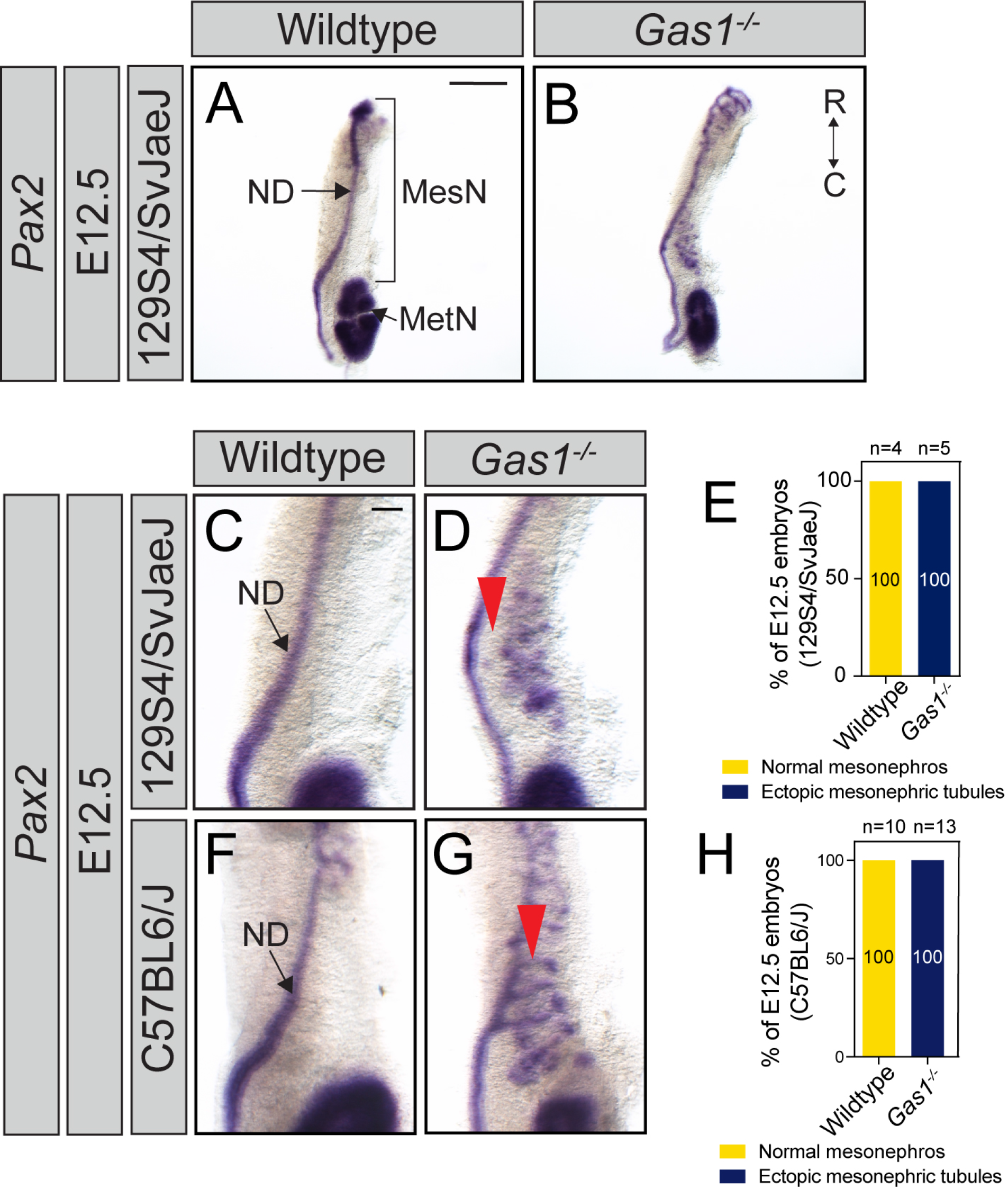
Ectopic caudal mesonephric tubules are maintained in *Gas1* mutant embryos on a congenic 129S4/SvJaeJ background. (A-B) Whole-mount *in situ* hybridization for *Pax2* in E12.5 wildtype (A) and *Gas1^-/-^* (B) mesonephros (MesN) and metanephros (MetN) from mice maintained on a congenic 129S4/SvJaeJ genetic background. ND, nephric duct; R, rostral; C, caudal. (C,D,F,G) Magnified images of whole-mount *in situ* hybridization for *Pax2* in E12.5 caudal mesonephros from wildtype (C) and *Gas1^-/-^*(D) embryos maintained on a 129S4/SvJaeJ background and wildtype (F) and *Gas1^-/-^*(G) embryos on a congenic C57BL/6J background. Red arrowheads identify regions of the caudal mesonephros in *Gas1* mutants that lack mesonephric tubule fusion to the ND (D, 129S4/SvJaeJ background) or are fused to the ND (G, C57BL6/J background). (E,H) Quantitation of the frequency of mesonephric defects in wildtype (n=4) and *Gas1^-/-^* (n=5) embryos maintained on a 129S4/SvJaeJ background or wildtype (n=10) and *Gas1^-/-^* (n=13) embryos maintained on a congenic C57BL6/J background classified into the following categories: normal mesonephros or ectopic caudal mesonephric tubules. Scale bar (A) 250μm; (C) 100μm. ND, nephric duct; R, rostral; C, caudal.

**Figure S5.**
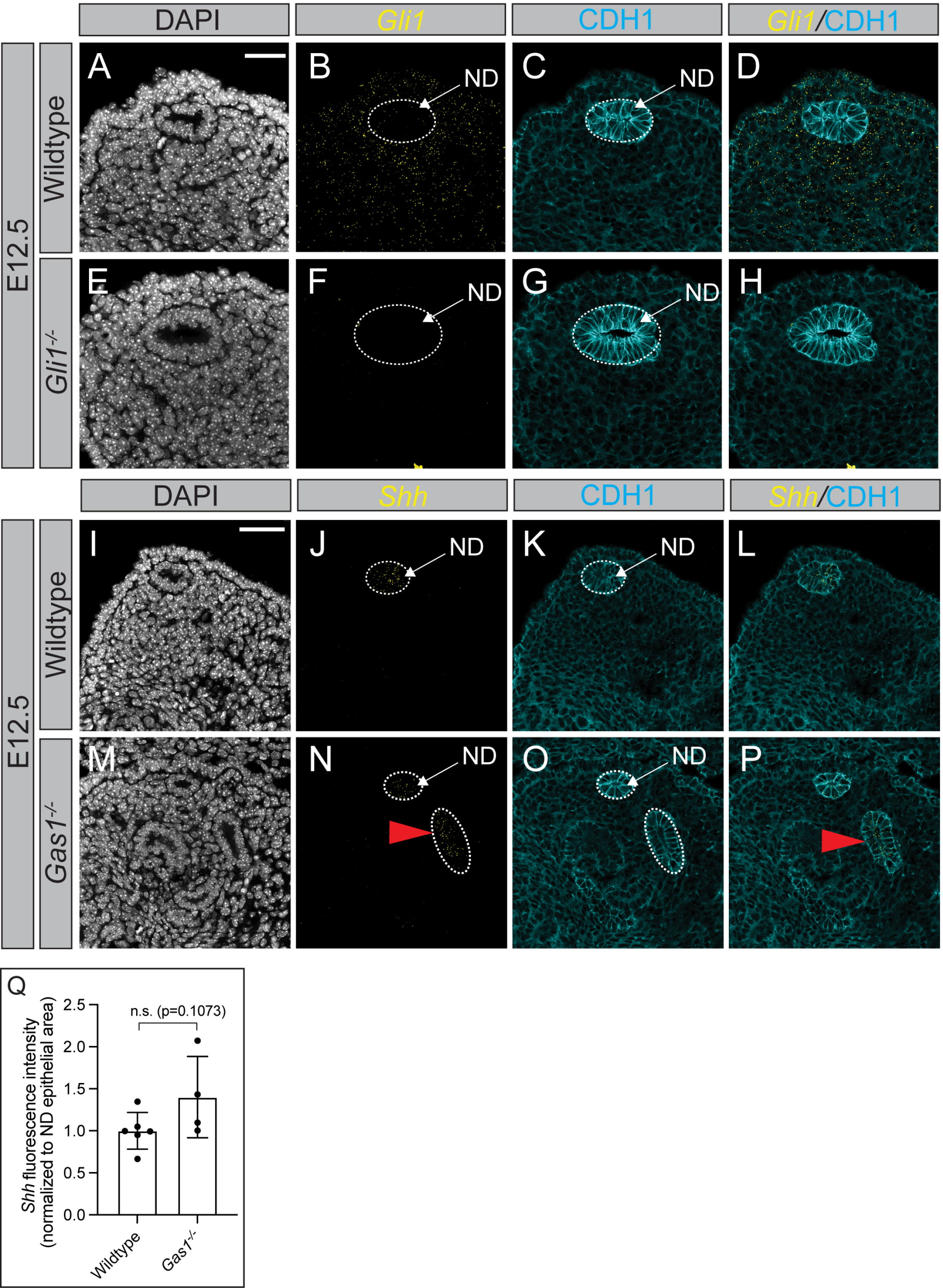
Ectopic *Shh* expression in *Gas1* mutant mesonephros. (A-H) FISH for *Gli1* (yellow; B,D,F,H) coupled with CDH1 antibody detection (cyan; C,D,G,H) in transverse caudal mesonephros sections from E12.5 wildtype (A-D) and *Gli1^-/-^* (E-H) embryos. DAPI labels nuclei (A,F). *Gli1*/CDH1 merged images are shown (D,H). Dotted lines mark the nephric duct (ND) epithelium (B,C,F,G). (I-P) FISH for *Shh* (yellow, J,N,L,P) coupled with CDH1 antibody detection (cyan; K,L,O,P) in transverse caudal mesonephros sections from E12.5 wildtype (I-L) and *Gas1^-/-^*(M-P) embryos. DAPI labels nuclei (I,M) *Shh/*CDH1 merged images are shown (L,P). Dotted lines mark the nephric duct (ND) epithelium (J,K,N,O) and ectopic *Shh-*expressing caudal mesonephric tubules in *Gas1* mutants (N,P; red arrowheads). (Q) Quantitation of *Shh* fluorescence intensity in transverse mesonephros sections from E12.5 wildtype (n=6 mesonephroi) and *Gas1^-/-^*(n=4 mesonephroi) normalized to ND area. For quantitation (Q), each data point (n) represents the average of at least 2 sections/kidney. Scale bars (A,I), 50μm. P-values were determined by a two-tailed Student’s t-test: non-significant: p>0.05 and significant: p≤0.05.

**Figure S6.**
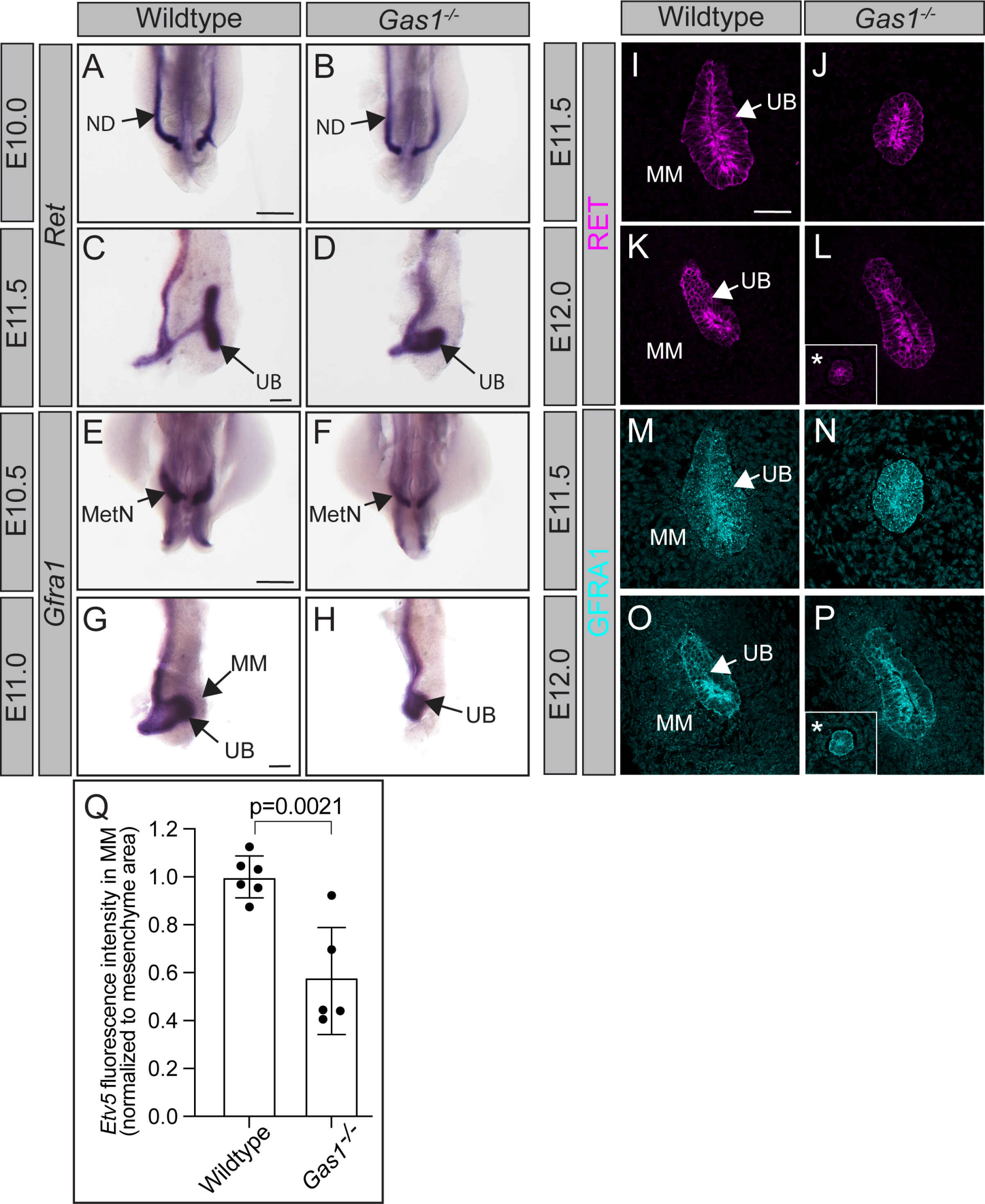
RET receptors are expressed normally in *Gas1^-/-^* metanephros. (A-H) Whole-mount *in situ* hybridization for *Ret* (A-D) and *Gfra1* (E-H) in wildtype (A,C,E,G) and *Gas1^-/-^* (B,D,F,H) metanephros (MetN). (I-P) Immunofluorescent antibody detection of RET (I-L) and GFRA1 (M-P) in coronal kidney sections from E11.5 and E12.0 wildtype (I,K,M,O) and *Gas1* mutant (J,L,N,P) embryos. Asterisks denote *Gas1* mutant kidneys that have not undergone ureteric bud (UB) branching morphogenesis (insets; L,P). (Q) Quantitation of *Etv5* MM fluorescence intensity normalized to MM area (3 sections/kidney) in E11.5 wildtype (n=6 kidneys) and *Gas1^-/-^* (n=5 kidneys) embryos. Scale bars (A,C,E,G) 100μm; (I) 50μm. P-values were determined by a two-tailed Student’s t-test: non-significant: p>0.05 and significant: p≤0.05.

**Figure S7.**
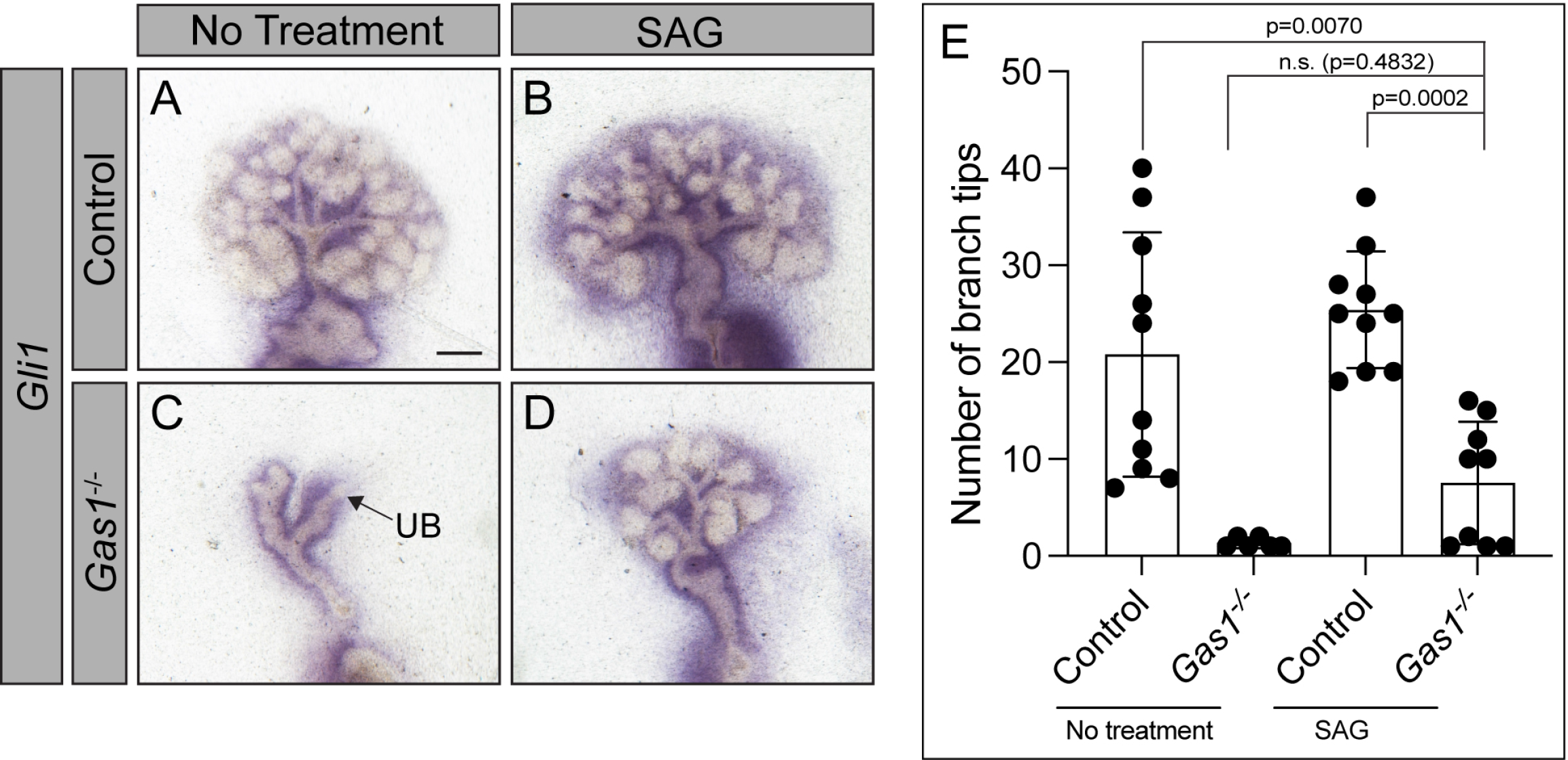
HH pathway activation does not rescue *Gas1^-/-^* branching defects. (A-D) Whole-mount *in situ* hybridization for *Gli1* expression in wildtype (A,B) and *Gas1^-/-^* (C,D) kidney explants cultured for 4 days in the presence or absence of 300nM Smoothened agonist (SAG). (E) Quantitation of the total number of UB branch tips in 4-day DMSO-treated or SAG-treated control (wildtype or *Gas1*^+/-^) and *Gas1^-/-^* kidney explant cultures. P-values were determined by an ordinary one-way ANOVA with Tukey’s multiple comparisons test: p>0.05 and significant: p≤0.05.

**Table S1.**
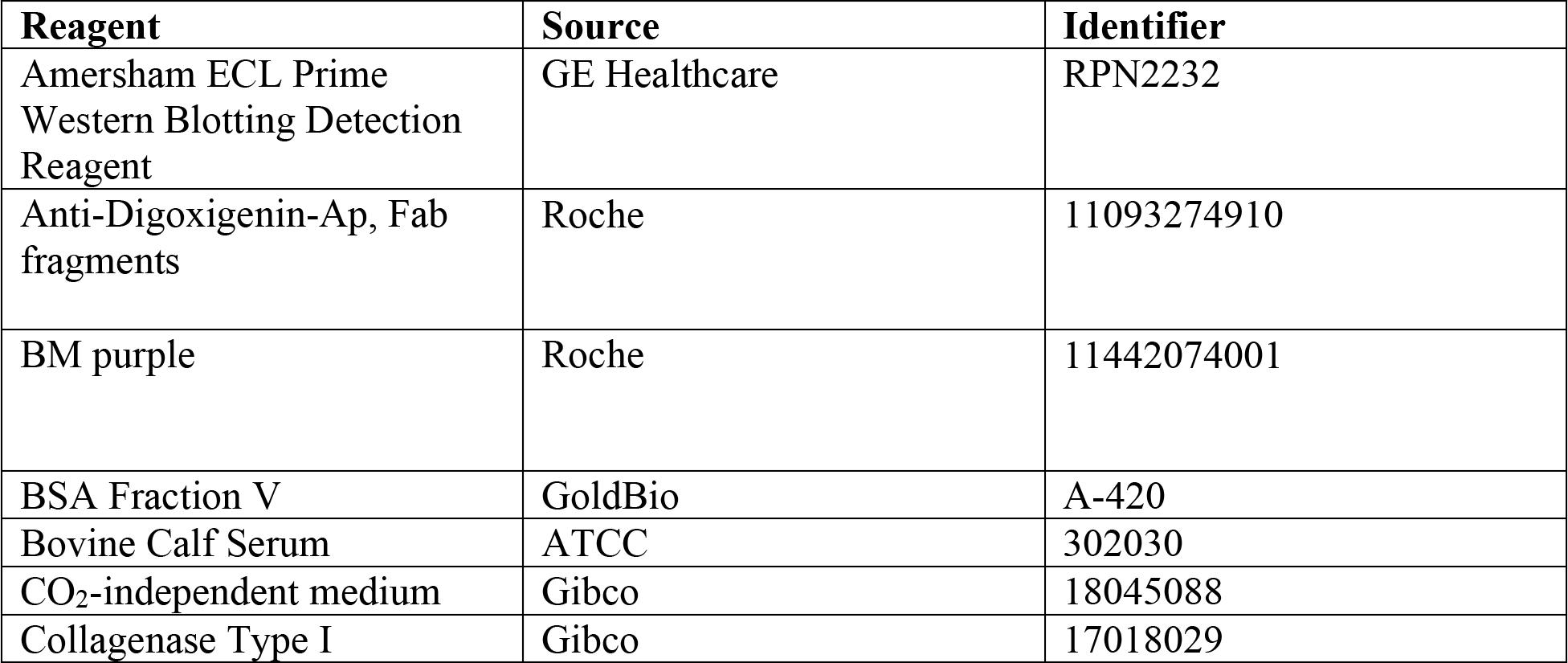
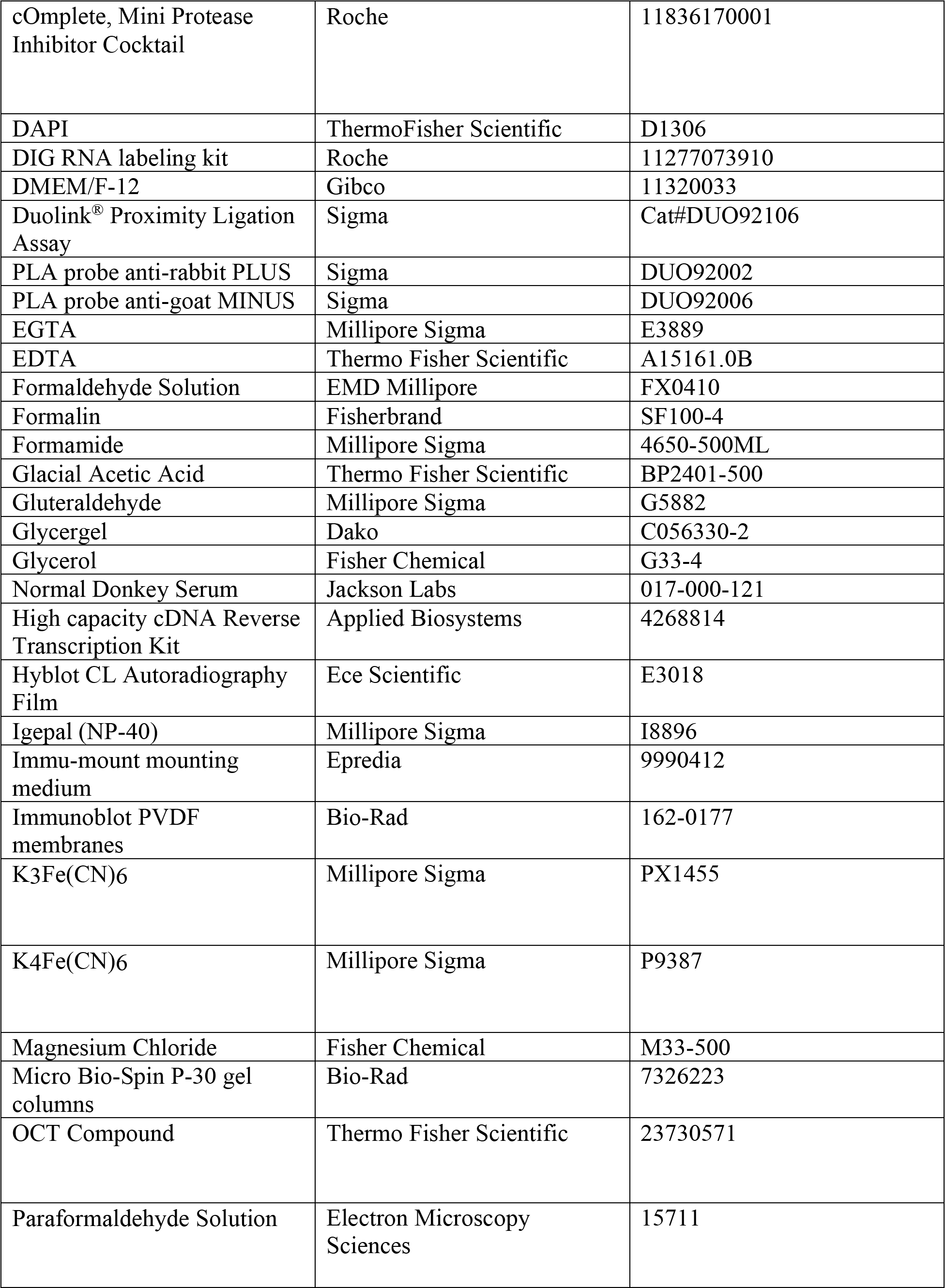
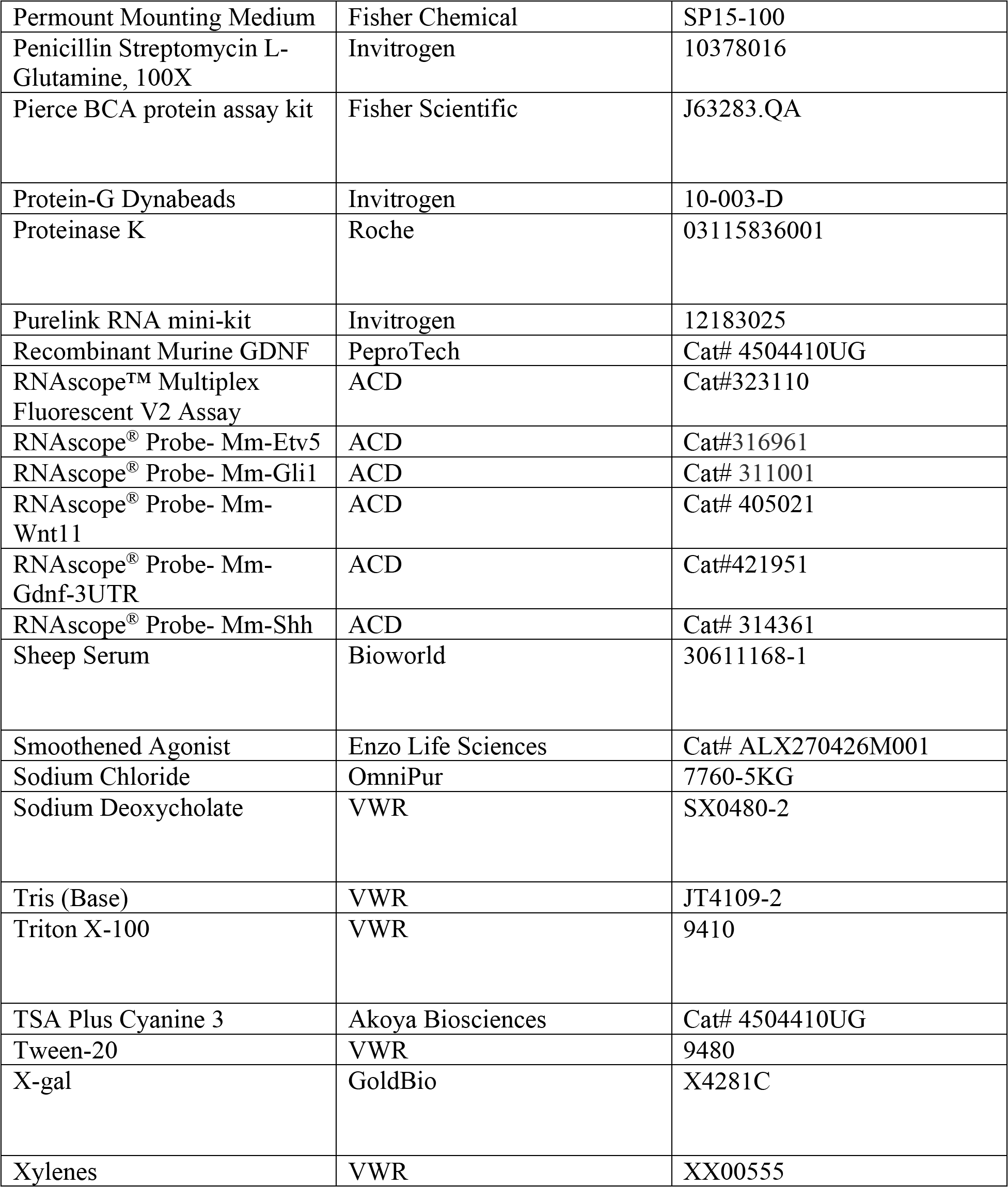
General Reagents.

**Table S2.** Primary Antibodies used for Immunofluorescence.

| Antibody | Source | Identifier | Dilution |
| --- | --- | --- | --- |
| Mouse monoclonal anti-Phospho-Histone H3 (SER10) (clone 6G3) | Cell Signaling Technology | Cat#9706S | 1:000 |
| Goat polyclonal anti-GAS1 | R&D Systems | Cat#AF2644;<br>RRID:AB_2107951 | 1:900 |
| Mouse monoclonal anti-E-Cadherin (CDH1) (clone 36) | BD Biosciences | Cat#610181;<br>RRID:AB_397581 | 1:500 |
| Rabbit monoclonal anti-Wilms Tumor 1 (clone CAN-R9(IHC)-56-2) | Abcam | Cat#ab89901;<br>RRID:AB_2043201 | 1:500 |
| Rabbit polyclonal anti-SIX2 | Proteintech | Cat#11562-1-AP;<br>RRID:AB_2189084 | 1:500 |
| Goat polyclonal anti-GFRA1 | R&D Systems | Cat#AF560;<br>RRID:AB_2110307 | 1:500 |
| Rabbit polyclonal anti-Cleaved Caspase-3 (Asp175) | Cell Signaling Technology | Cat#9661;<br>RRID:AB_2341188 | 1:200 |
| Sheep polyclonal anti-Digoxigenin-AP, Fab fragments | Roche | Cat#11093274910;<br>RRID:AB_514497 | 1:4000 |
| Rat monoclonal anti-HSPG2 (Perlecan) (Clone A7L6) | Millipore | Cat#MAB1948P;<br>RRID:AB_10615958 | 1:500 |
| Chicken polyclonal anti-Beta-Galactosidase | Abcam | Cat#ab9361;<br>RRID:AB_307210 | 1:2500 |
| Rabbit polyclonal anti-PAX2 | Thermo Fisher Scientific | Cat#71-6000;<br>RRID:AB_2533990 | 1:250 |
| Rabbit monoclonal anti-RET (Clone C31B4) | Cell Signaling Technology | Cat#3223;<br>RRID:AB_2238465 | 1:200 |

**Table S3.** Experimental mouse models.

| Strain | Source | Identifier |
| --- | --- | --- |
| Mouse:*Gas1<sup>lacZ</sup>* | (Martinelli and Fan, 2007a) | N/A |
| Mouse:129S4/SvJaeJ | The Jackson Laboratory | RRID:IMSR_JAX:009104 |
| Mouse:C57BL6/J | The Jackson Laboratory | RRID:IMSR_JAX:000664 |
| Mouse:*Boc<sup>AP</sup>* | (Zhang et al., 2011) | N/A |
| Mouse:*Cdon*<sup>lacZ-1</sup> | (Cole and Krauss, 2003) | N/A |
| Mouse:*Cdon*<sup>lacZ-2</sup> | (Cole and Krauss, 2003) | N/A |
| Mouse:129S/Sv-*Ret*<sup>tm1Cos/J</sup> | (Schuchardt et al., 1994) | RRID:IMSR_JAX:009085 |

## References

1. Allen, B.L., Song, J.Y., Izzi, L., Althaus, I.W., Kang, J.S., Charron, F., Krauss, R.S., McMahon, A.P., 2011. Overlapping roles and collective requirement for the coreceptors GAS1, CDO, and BOC in SHH pathway function. Dev Cell 20, 775–787.

2. Allen, B.L., Tenzen, T., McMahon, A.P., 2007. The Hedgehog-binding proteins Gas1 and Cdo cooperate to positively regulate Shh signaling during mouse development. Genes Dev 21, 1244–1257.

3. Bai, C.B., Joyner, A.L., 2001. Gli1 can rescue the in vivo function of Gli2. Development 128, 5161–5172.

4. Barak, H., Boyle, S.C., 2011. Organ culture and immunostaining of mouse embryonic kidneys. Cold Spring Harb Protoc 2011, pdb prot5558.

5. Biau, S., Jin, S., Fan, C.M., 2013. Gastrointestinal defects of the Gas1 mutant involve dysregulated Hedgehog and Ret signaling. Biol Open 2, 144–155.

6. Bohnenpoll, T., Wittern, A.B., Mamo, T.M., Weiss, A.C., Rudat, C., Kleppa, M.J., Schuster- Gossler, K., Wojahn, I., Ludtke, T.H., Trowe, M.O., Kispert, A., 2017. A SHH-FOXF1-BMP4 signaling axis regulating growth and differentiation of epithelial and mesenchymal tissues in ureter development. PLoS Genet 13, e1006951.

7. Cabrera, J.R., Sanchez-Pulido, L., Rojas, A.M., Valencia, A., Manes, S., Naranjo, J.R., Mellstrom, B., 2006. Gas1 is related to the glial cell-derived neurotrophic factor family receptors alpha and regulates Ret signaling. J Biol Chem 281, 14330–14339.

8. Cacalano, G., Farinas, I., Wang, L.C., Hagler, K., Forgie, A., Moore, M., Armanini, M., Phillips, H., Ryan, A.M., Reichardt, L.F., Hynes, M., Davies, A., Rosenthal, A., 1998. GFRalpha1 is an essential receptor component for GDNF in the developing nervous system and kidney. Neuron 21, 53–62.

9. Cain, J.E., Islam, E., Haxho, F., Chen, L., Bridgewater, D., Nieuwenhuis, E., Hui, C.C., Rosenblum, N.D., 2009. GLI3 repressor controls nephron number via regulation of Wnt11 and Ret in ureteric tip cells. PLoS One 4, e7313.

10. Chu, J., Ding, J., Jeays-Ward, K., Price, S.M., Placzek, M., Shen, M.M., 2005. Non-cell- autonomous role for Cripto in axial midline formation during vertebrate embryogenesis. Development 132, 5539–5551.

11. Chung, E., Deacon, P., Hu, Y.C., Lim, H.W., Park, J.S., 2024. Hedgehog signaling is required for the maintenance of mesenchymal nephron progenitors. bioRxiv.

12. Cobourne, M.T., Miletich, I., Sharpe, P.T., 2004. Restriction of sonic hedgehog signalling during early tooth development. Development 131, 2875–2885.

13. Cole, F., Krauss, R.S., 2003. Microform holoprosencephaly in mice that lack the Ig superfamily member Cdon. Curr Biol 13, 411–415.

14. Combes, A.N., Davies, J.A., Little, M.H., 2015. Cell-cell interactions driving kidney morphogenesis. Curr Top Dev Biol 112, 467–508.

15. Costantini, F., Kopan, R., 2010. Patterning a complex organ: branching morphogenesis and nephron segmentation in kidney development. Dev Cell 18, 698–712.

16. Costantini, F., Watanabe, T., Lu, B., Chi, X., Srinivas, S., 2011. Dissection of embryonic mouse kidney, culture in vitro, and imaging of the developing organ. Cold Spring Harb Protoc 2011, pdb prot5613.

17. Creanga, A., Glenn, T.D., Mann, R.K., Saunders, A.M., Talbot, W.S., Beachy, P.A., 2012. Scube/You activity mediates release of dually lipid-modified Hedgehog signal in soluble form. Genes Dev 26, 1312–1325.

18. Dai, P., Akimaru, H., Tanaka, Y., Maekawa, T., Nakafuku, M., Ishii, S., 1999. Sonic Hedgehog- induced activation of the Gli1 promoter is mediated by GLI3. J Biol Chem 274, 8143–8152.

19. Dobrowolski, M., Cave, C., Levy-Myers, R., Lee, C., Park, S., Choi, B.R., Xiao, B., Yang, W., Sockanathan, S., 2020. GDE3 regulates oligodendrocyte precursor proliferation via release of soluble CNTFRalpha. Development 147.

20. Dressler, G.R., Deutsch, U., Chowdhury, K., Nornes, H.O., Gruss, P., 1990. Pax2, a new murine paired-box-containing gene and its expression in the developing excretory system. Development 109, 787–795.

21. Durbec, P., Marcos-Gutierrez, C.V., Kilkenny, C., Grigoriou, M., Wartiowaara, K., Suvanto, P., Smith, D., Ponder, B., Costantini, F., Saarma, M., et al., 1996. GDNF signalling through the Ret receptor tyrosine kinase. Nature 381, 789–793.

22. Echevarria-Andino, M.L., Allen, B.L., 2020. The hedgehog co-receptor BOC differentially regulates SHH signaling during craniofacial development. Development 147.

23. Echevarria-Andino, M.L., Franks, N.E., Schrader, H.E., Hong, M., Krauss, R.S., Allen, B.L., 2022. CDON contributes to Hedgehog-dependent patterning and growth of the developing limb. Dev Biol 493, 1–11.

24. Enomoto, H., Hughes, I., Golden, J., Baloh, R.H., Yonemura, S., Heuckeroth, R.O., Johnson, E.M., Jr., Milbrandt, J., 2004. GFRalpha1 expression in cells lacking RET is dispensable for organogenesis and nerve regeneration. Neuron 44, 623–636.

25. Fleming, M.S., Vysochan, A., Paixao, S., Niu, J., Klein, R., Savitt, J.M., Luo, W., 2015. Cis and trans RET signaling control the survival and central projection growth of rapidly adapting mechanoreceptors. Elife 4, e06828.

26. Grieshammer, U., Le, M., Plump, A.S., Wang, F., Tessier-Lavigne, M., Martin, G.R., 2004. SLIT2-mediated ROBO2 signaling restricts kidney induction to a single site. Dev Cell 6, 709–717.

27. Holtz, A.M., Griffiths, S.C., Davis, S.J., Bishop, B., Siebold, C., Allen, B.L., 2015. Secreted HHIP1 interacts with heparan sulfate and regulates Hedgehog ligand localization and function. J Cell Biol 209, 739–757.

28. Huang, P., Wierbowski, B.M., Lian, T., Chan, C., Garcia-Linares, S., Jiang, J., Salic, A., 2022. Structural basis for catalyzed assembly of the Sonic hedgehog-Patched1 signaling complex. Dev Cell 57, 670–685 e678.

29. Izzi, L., Levesque, M., Morin, S., Laniel, D., Wilkes, B.C., Mille, F., Krauss, R.S., McMahon, A.P., Allen, B.L., Charron, F., 2011. Boc and Gas1 each form distinct Shh receptor complexes with Ptch1 and are required for Shh-mediated cell proliferation. Dev Cell 20, 788–801.

30. Jin, S., Martinelli, D.C., Zheng, X., Tessier-Lavigne, M., Fan, C.M., 2015. Gas1 is a receptor for sonic hedgehog to repel enteric axons. Proc Natl Acad Sci U S A 112, E73–80.

31. Jing, S., Wen, D., Yu, Y., Holst, P.L., Luo, Y., Fang, M., Tamir, R., Antonio, L., Hu, Z., Cupples, R., Louis, J.C., Hu, S., Altrock, B.W., Fox, G.M., 1996. GDNF-induced activation of the ret protein tyrosine kinase is mediated by GDNFR-alpha, a novel receptor for GDNF. Cell 85, 1113–1124.

32. Kang, J.S., Mulieri, P.J., Hu, Y., Taliana, L., Krauss, R.S., 2002. BOC, an Ig superfamily member, associates with CDO to positively regulate myogenic differentiation. EMBO J 21, 114–124.

33. Kann, M., Bae, E., Lenz, M.O., Li, L., Trannguyen, B., Schumacher, V.A., Taglienti, M.E., Bordeianou, L., Hartwig, S., Rinschen, M.M., Schermer, B., Benzing, T., Fan, C.M., Kreidberg, J.A., 2015. WT1 targets Gas1 to maintain nephron progenitor cells by modulating FGF signals. Development 142, 1254–1266.

34. Kume, T., Deng, K., Hogan, B.L., 2000. Murine forkhead/winged helix genes Foxc1 (Mf1) and Foxc2 (Mfh1) are required for the early organogenesis of the kidney and urinary tract. Development 127, 1387–1395.

35. Ledda, F., Paratcha, G., Ibanez, C.F., 2002. Target-derived GFRalpha1 as an attractive guidance signal for developing sensory and sympathetic axons via activation of Cdk5. Neuron 36, 387–401.

36. Lee, C.S., Buttitta, L., Fan, C.M., 2001. Evidence that the WNT-inducible growth arrest-specific gene 1 encodes an antagonist of sonic hedgehog signaling in the somite. Proc Natl Acad Sci U S A 98, 11347–11352.

37. Lee, C.S., Fan, C.M., 2001. Embryonic expression patterns of the mouse and chick Gas1 genes. Mech Dev 101, 293–297.

38. Lee, G.H., Fujita, M., Takaoka, K., Murakami, Y., Fujihara, Y., Kanzawa, N., Murakami, K.I., Kajikawa, E., Takada, Y., Saito, K., Ikawa, M., Hamada, H., Maeda, Y., Kinoshita, T., 2016. A GPI processing phospholipase A2, PGAP6, modulates Nodal signaling in embryos by shedding CRIPTO. J Cell Biol 215, 705-718.

39. Li, L., Rozo, M., Yue, S., Zheng, X., F, J.T., Lepper, C., Fan, C.M., 2019. Muscle stem cell renewal suppressed by Gas1 can be reversed by GDNF in mice. Nat Metab 1, 985–995.

40. Little, M.H., McMahon, A.P., 2012. Mammalian kidney development: principles, progress, and projections. Cold Spring Harb Perspect Biol 4.

41. Liu, Y., May, N.R., Fan, C.M., 2001. Growth arrest specific gene 1 is a positive growth regulator for the cerebellum. Dev Biol 236, 30–45.

42. Lopez-Ramirez, M.A., Dominguez-Monzon, G., Vergara, P., Segovia, J., 2008. Gas1 reduces Ret tyrosine 1062 phosphorylation and alters GDNF-mediated intracellular signaling. Int J Dev Neurosci 26, 497–503.

43. Lu, B.C., Cebrian, C., Chi, X., Kuure, S., Kuo, R., Bates, C.M., Arber, S., Hassell, J., MacNeil, L., Hoshi, M., Jain, S., Asai, N., Takahashi, M., Schmidt-Ott, K.M., Barasch, J., D’Agati, V., Costantini, F., 2009. Etv4 and Etv5 are required downstream of GDNF and Ret for kidney branching morphogenesis. Nat Genet 41, 1295–1302.

44. Majumdar, A., Vainio, S., Kispert, A., McMahon, J., McMahon, A.P., 2003. Wnt11 and Ret/Gdnf pathways cooperate in regulating ureteric branching during metanephric kidney development. Development 130, 3175–3185.

45. Marczenke, M., Sunaga-Franze, D.Y., Popp, O., Althaus, I.W., Sauer, S., Mertins, P., Christ, A., Allen, B.L., Willnow, T.E., 2021. GAS1 is required for NOTCH-dependent facilitation of SHH signaling in the ventral forebrain neuroepithelium. Development 148.

46. Martinelli, D.C., Fan, C.M., 2007a. Gas1 extends the range of Hedgehog action by facilitating its signaling. Genes Dev 21, 1231–1243.

47. Martinelli, D.C., Fan, C.M., 2007b. The role of Gas1 in embryonic development and its implications for human disease. Cell Cycle 6, 2650–2655.

48. McKean, M., Napoli, F.R., Hasan, T., Joseph, T., Wheeler, A., Beebe, K., Soriano-Cruz, S., Kawano, M., Cave, C., 2023. GDE6 promotes progenitor identity in the vertebrate neural tube. Front Neurosci 17, 1047767.

49. McMahon, A.P., Aronow, B.J., Davidson, D.R., Davies, J.A., Gaido, K.W., Grimmond, S., Lessard, J.L., Little, M.H., Potter, S.S., Wilder, E.L., Zhang, P., project, G., 2008. GUDMAP: the genitourinary developmental molecular anatomy project. J Am Soc Nephrol 19, 667–671.

50. Moore, M.W., Klein, R.D., Farinas, I., Sauer, H., Armanini, M., Phillips, H., Reichardt, L.F., Ryan, A.M., Carver-Moore, K., Rosenthal, A., 1996. Renal and neuronal abnormalities in mice lacking GDNF. Nature 382, 76-79.

51. Mulieri, P.J., Kang, J.S., Sassoon, D.A., Krauss, R.S., 2002. Expression of the boc gene during murine embryogenesis. Dev Dyn 223, 379–388.

52. Mulieri, P.J., Okada, A., Sassoon, D.A., McConnell, S.K., Krauss, R.S., 2000. Developmental expression pattern of the cdo gene. Dev Dyn 219, 40–49.

53. Muraguchi, T., Takegami, Y., Ohtsuka, T., Kitajima, S., Chandana, E.P., Omura, A., Miki, T., Takahashi, R., Matsumoto, N., Ludwig, A., Noda, M., Takahashi, C., 2007. RECK modulates Notch signaling during cortical neurogenesis by regulating ADAM10 activity. Nat Neurosci 10, 838–845.

54. Murashima, A., Akita, H., Okazawa, M., Kishigami, S., Nakagata, N., Nishinakamura, R., Yamada, G., 2014. Midline-derived Shh regulates mesonephric tubule formation through the paraxial mesoderm. Dev Biol 386, 216–226.

55. Ohazama, A., Haycraft, C.J., Seppala, M., Blackburn, J., Ghafoor, S., Cobourne, M., Martinelli, D.C., Fan, C.M., Peterkova, R., Lesot, H., Yoder, B.K., Sharpe, P.T., 2009. Primary cilia regulate Shh activity in the control of molar tooth number. Development 136, 897–903.

56. Paratcha, G., Ledda, F., Baars, L., Coulpier, M., Besset, V., Anders, J., Scott, R., Ibanez, C.F., 2001. Released GFRalpha1 potentiates downstream signaling, neuronal survival, and differentiation via a novel mechanism of recruitment of c-Ret to lipid rafts. Neuron 29, 171–184.

57. Parisi, S., D’Andrea, D., Lago, C.T., Adamson, E.D., Persico, M.G., Minchiotti, G., 2003. Nodal- dependent Cripto signaling promotes cardiomyogenesis and redirects the neural fate of embryonic stem cells. J Cell Biol 163, 303-314.

58. Park, S., Lee, C., Sabharwal, P., Zhang, M., Meyers, C.L., Sockanathan, S., 2013. GDE2 promotes neurogenesis by glycosylphosphatidylinositol-anchor cleavage of RECK. Science 339, 324–328.

59. Pichel, J.G., Shen, L., Sheng, H.Z., Granholm, A.C., Drago, J., Grinberg, A., Lee, E.J., Huang, S.P., Saarma, M., Hoffer, B.J., Sariola, H., Westphal, H., 1996a. Defects in enteric innervation and kidney development in mice lacking GDNF. Nature 382, 73–76.

60. Pichel, J.G., Shen, L., Sheng, H.Z., Granholm, A.C., Drago, J., Grinberg, A., Lee, E.J., Huang, S.P., Saarma, M., Hoffer, B.J., Sariola, H., Westphal, H., 1996b. GDNF is required for kidney development and enteric innervation. Cold Spring Harb Symp Quant Biol 61, 445–457.

61. Rosti, K., Goldman, A., Kajander, T., 2015. Solution structure and biophysical characterization of the multifaceted signalling effector protein growth arrest specific-1. BMC Biochem 16, 8.

62. Rowan, C.J., Li, W., Martirosyan, H., Erwood, S., Hu, D., Kim, Y.K., Sheybani-Deloui, S., Mulder, J., Blake, J., Chen, L., Rosenblum, N.D., 2018. Hedgehog-GLI signaling in Foxd1- positive stromal cells promotes murine nephrogenesis via TGFbeta signaling. Development 145.

63. Sabharwal, P., Lee, C., Park, S., Rao, M., Sockanathan, S., 2011. GDE2 regulates subtype- specific motor neuron generation through inhibition of Notch signaling. Neuron 71, 1058-1070.

64. Sanchez, M.P., Silos-Santiago, I., Frisen, J., He, B., Lira, S.A., Barbacid, M., 1996. Renal agenesis and the absence of enteric neurons in mice lacking GDNF. Nature 382, 70-73.

65. Schneider, C., King, R.M., Philipson, L., 1988. Genes specifically expressed at growth arrest of mammalian cells. Cell 54, 787–793.

66. Schneider, C.A., Rasband, W.S., Eliceiri, K.W., 2012. NIH Image to ImageJ: 25 years of image analysis. Nat Methods 9, 671–675.

67. Schuchardt, A., D’Agati, V., Larsson-Blomberg, L., Costantini, F., Pachnis, V., 1994. Defects in the kidney and enteric nervous system of mice lacking the tyrosine kinase receptor Ret. Nature 367, 380–383.

68. Schuchardt, A., D’Agati, V., Pachnis, V., Costantini, F., 1996. Renal agenesis and hypodysplasia in ret-k- mutant mice result from defects in ureteric bud development. Development 122, 1919–1929.

69. Seppala, M., Depew, M.J., Martinelli, D.C., Fan, C.M., Sharpe, P.T., Cobourne, M.T., 2007. Gas1 is a modifier for holoprosencephaly and genetically interacts with sonic hedgehog. J Clin Invest 117, 1575–1584.

70. Seppala, M., Thivichon-Prince, B., Xavier, G.M., Shaffie, N., Sangani, I., Birjandi, A.A., Rooney, J., Lau, J.N.S., Dhaliwal, R., Rossi, O., Riaz, M.A., Stonehouse-Smith, D., Wang, Y., Papageorgiou, S.N., Viriot, L., Cobourne, M.T., 2022. Gas1 Regulates Patterning of the Murine and Human Dentitions through Sonic Hedgehog. J Dent Res 101, 473–482.

71. Seppala, M., Xavier, G.M., Fan, C.M., Cobourne, M.T., 2014. Boc modifies the spectrum of holoprosencephaly in the absence of Gas1 function. Biol Open 3, 728–740.

72. Song, J.Y., Holtz, A.M., Pinskey, J.M., Allen, B.L., 2015. Distinct structural requirements for CDON and BOC in the promotion of Hedgehog signaling. Dev Biol 402, 239–252.

73. Treanor, J.J., Goodman, L., de Sauvage, F., Stone, D.M., Poulsen, K.T., Beck, C.D., Gray, C., Armanini, M.P., Pollock, R.A., Hefti, F., Phillips, H.S., Goddard, A., Moore, M.W., Buj-Bello, A., Davies, A.M., Asai, N., Takahashi, M., Vandlen, R., Henderson, C.E., Rosenthal, A., 1996. Characterization of a multicomponent receptor for GDNF. Nature 382, 80–83.

74. Trupp, M., Arenas, E., Fainzilber, M., Nilsson, A.S., Sieber, B.A., Grigoriou, M., Kilkenny, C., Salazar-Grueso, E., Pachnis, V., Arumae, U., 1996. Functional receptor for GDNF encoded by the c-ret proto-oncogene. Nature 381, 785–789.

75. Tukachinsky, H., Kuzmickas, R.P., Jao, C.Y., Liu, J., Salic, A., 2012. Dispatched and scube mediate the efficient secretion of the cholesterol-modified hedgehog ligand. Cell Rep 2, 308–320.

76. van Roeyen, C.R., Zok, S., Pruessmeyer, J., Boor, P., Nagayama, Y., Fleckenstein, S., Cohen, C.D., Eitner, F., Grone, H.J., Ostendorf, T., Ludwig, A., Floege, J., 2013. Growth arrest-specific protein 1 is a novel endogenous inhibitor of glomerular cell activation and proliferation. Kidney Int 83, 251-263.

77. Watanabe, K., Hamada, S., Bianco, C., Mancino, M., Nagaoka, T., Gonzales, M., Bailly, V., Strizzi, L., Salomon, D.S., 2007. Requirement of glycosylphosphatidylinositol anchor of Cripto- 1 for trans activity as a Nodal co-receptor. J Biol Chem 282, 35772–35786.

78. Yan, Y.T., Liu, J.J., Luo, Y., E, C., Haltiwanger, R.S., Abate-Shen, C., Shen, M.M., 2002. Dual roles of Cripto as a ligand and coreceptor in the nodal signaling pathway. Mol Cell Biol 22, 4439–4449.

79. Yu, J., Carroll, T.J., McMahon, A.P., 2002. Sonic hedgehog regulates proliferation and differentiation of mesenchymal cells in the mouse metanephric kidney. Development 129, 5301–5312.

80. Yu, T., Scully, S., Yu, Y., Fox, G.M., Jing, S., Zhou, R., 1998. Expression of GDNF family receptor components during development: implications in the mechanisms of interaction. J Neurosci 18, 4684–4696.

81. Zhang, W., Hong, M., Bae, G.U., Kang, J.S., Krauss, R.S., 2011. Boc modifies the holoprosencephaly spectrum of Cdo mutant mice. Dis Model Mech 4, 368–380.

82. Zhou, B., Ma, Q., Rajagopal, S., Wu, S.M., Domian, I., Rivera-Feliciano, J., Jiang, D., von Gise, A., Ikeda, S., Chien, K.R., Pu, W.T., 2008. Epicardial progenitors contribute to the cardiomyocyte lineage in the developing heart. Nature 454, 109–113.

